# Towards a mechanistic understanding of collective escape in starlings

**DOI:** 10.1101/2024.10.27.620514

**Authors:** Marina Papadopoulou, Hanno Hildenbrandt, Rolf F. Storms, Claudio Carere, Simon Verhulst, Charlotte K. Hemelrijk

## Abstract

Large flocks of European starlings change shape, size, and internal structure continuously and rapidly when hunted by aerial predators. How their diverse patterns of collective escape emerge is still unknown. Here, we disentangle the collective behavior of starlings combining video footage of flocks pursued by a robotic predator and a data-driven 3-dimentional agent-based model. In vivo, we show that flock members often differ in their evasive maneuvers and that several collective patterns arise simultaneously across a flock. In silica, we identify individual-level rules that lead to dynamics of collective motion and escape similar to real starlings. Our results suggest that the mechanisms underlying starling murmurations depend on how fast escape information propagates through the flock, the relative positions of the escaping individuals to the predator, and the previous state of the flock (hysteresis). Our study highlights the importance of investigating fine-scale dynamics and micro-macro feedback when studying self-organized adaptive systems.

## Introduction

Some of the most complex patterns of collective behavior in animals are displayed when groups are threatened by a predator. Rapid changes in shape and internal structure of the group confuse the predator ^1,2^ and reduce the flock members’ risk of getting caught ^3,4^. The murmurations of European starlings (*Sturnus vulgaris*) ^5^ are one of the best-known phenomena of collective behavior ^5,6^; their large flocks exhibit a great diversity of complex patterns, in particular when attacked by aerial predators such as the peregrine falcon (*Falco peregrinus*) ^7,8^. However, besides insight into the collective motion of their flocks ^9–12^, little is known about what rules of motion and interaction underlie their patterns of collective escape ^13,14^, knowledge that is vital to understand the adaptive value of these social interactions and the formation of the spatiotemporal patterns we observe in nature.

In starlings, many patterns of collective escape can be distinguished based on shape, darkness, size, internal structure and dynamics of the flock (similar to other systems^15–18^), namely the agitation wave, the vacuole, the flash expansion, the collective turn, the cordon, the split, blackening, compacting, and dilution ^8^. During an agitation wave, a dark band moves from one side of the flock (closer to the predator) to the other ^8,19^. During flash expansion, flock members suddenly move radially outwards (usually away from the attacking predator) ^8,18^. The split implies that the flock divides into two or more parts, referred to as sub-flocks ^8,20^. During blackening, the flock (or part thereof) appears darker, and during compacting smaller^8^. During dilution, the distance between flock members increases and the flock becomes lighter in color^8^. The vacuole refers to a hole in a polarized flock with the individuals around the hole moving in the same direction, and the cordon to the thin line (cordon) that connects two relatively large parts of the flock ^7,8^. Earlier studies have only focused on the relation between the frequency of these patterns and the frequency and timing of predator attacks ^7,8,19^, without investigating the mechanisms driving their emergence.

The link between individual behavior and global patterns can be detected through computational models based on self-organization although it is important to note that in a simulation, the same macroscopic patterns may emerge from different rules of individual behavior. There is thus no guarantee that models represent correctly individual behavior of animals in nature but they offer a theoretical understanding of what is possible given a set of assumptions. To increase the biological relevance of the theoretical predictions of a model, the local interactions and individual motion of simulated agents can be adjusted to empirical data^21–23^. Model predictions can then in turn drive the collection of more empirical data in order to validate or challenge our theoretical understanding^23^. In the case of collective motion, models usually include moving agents that align with and are attracted to each other while avoiding collisions ^24–26^ (but see also ^27^). In these models, the emergence of several collective properties have been examined such as flock diffusion ^28,29^, milling ^26,30^, and flock shape ^21,31^.

In the case of collective escape, the development of computational models of bird flocks has been constrained by the difficulty of empirical data collection^10,11,13,20,32,33^. So far, only a few patterns of collective escape of starlings and of homing pigeons (*Columba livia*) have been studied in detail ^13,14,20,32^, using the computational models StarDisplay ^11^ and HoPE ^32^, respectively. Agitation waves in simulated flocks of starlings were shown to emerge from a fixed zig-zag motion performed by a few individuals close to the predator and copied by their neighbors ^13^. Splits and collective turns in pigeons were shown to emerge from a single individual escape maneuver involving discrete, sudden turns interrupting the continuous coordinated motion of the group ^20^. Mathematical models have focused on the temporal link of pairs of patterns, showing how splits may follow flash expansions and agitation waves blackening^34^. Mechanistical insight about the spatiotemporal link between patterns is rare in the literature; most computational models have focused on the emergence of a single type of collective escape (but see ^16,35,36^ for studies on fish schools). The simultaneous occurrence of several patterns of collective escape, like in the large murmurations of starlings in particular, have to our best knowledge not been investigated previously.

In the present paper, we examine the mechanisms underlying several patterns of collective escape in starlings, bridging the gap between empirical observations and theory. We analyzed video footage assembled in the field by employing a bio-hybrid solution that overcomes the unpredictability of predator attacks in nature ^37,38^: flocks of starlings under attack by a remotely-controlled robotic predator, the RobotFalcon ^39^. Previous work has established that flocks respond to the RobotFalcon similarly to the real peregrine falcon ^39^. Our empirical investigation showed that patterns of collective escape often arise simultaneously in a single flock. To study the emergence of this phenomenon, we developed a biologically-inspired computational model of collective motion and escape in 3-dimensions specifically for starlings, through pattern-oriented modelling ^22^ and the use of non-homogenous Markov-chains at the individual level decision making. We simulated flocks of up to a few thousands of individuals under attack and identified the conditions at which different patterns emerge. We further discovered that hysteresis (collective memory) may underlie the density and shape dynamics of a flock, i.e., its current state is affected by its previous state ^26,40^.

## Results

### Collective escape of starling flocks

We analyzed data^39,41^ in which the RobotFalcon pursued starling flocks of about 20 to 2000 individuals for approximately 20 seconds to 2 minutes (with average pursuit duration of 1 min). We focused our analysis on the flocks in our dataset in which more than one pattern of collective escape co-occurred (n=10). Video footage of the pursuit was recorded by both a ground camera and a camera on the RobotFalcon (see supplementary Fig.S1 for both camera views)^39^.

The patterns of collective escape we observed included the collective turn (Fig. 1A), the split (supplementary Fig. S2), the cordon, the flash expansion, the agitation wave, blackening, compacting (supplementary Fig. S3) and dilution. Starlings also executed ‘columnar flocking’: their flocks often took a vertical shape, elongated in altitude (Fig. 1B, supplementary Movie S1, S2B), something previously characterized only in flocks of dunlins (*Calidris alpina*) ^42,43^. Multiple patterns of collective escape sometimes arose simultaneously in a single flock (Fig. 1B). Specifically the collective turn most often co-occurred with the split, the columnar flocking, or the compacting (Fig. 1C). The cordon seems to arise as a result of delayed information transfer during columnar flocking, when the top of the flock performed consecutive collective turns and the bottom part was following with a small delay (Fig. 1D). The flash expansion appeared at the point where the RobotFalcon attacked (most often in the periphery of the flock), while at the same time in another part of the flock a collective turn or compacting was taking place. Agitation waves were observed only in the largest flock in our dataset. Since the emergence of the agitation wave and the flash expansion have been studied previously ^8,13,14,18^, we focused our analysis on the other patterns (see supplementary Movie S1). We present a simplified schematic of part of this escape sequence in Fig. 1D, showing that parts of the flock behave differently.

**Fig. 1.**
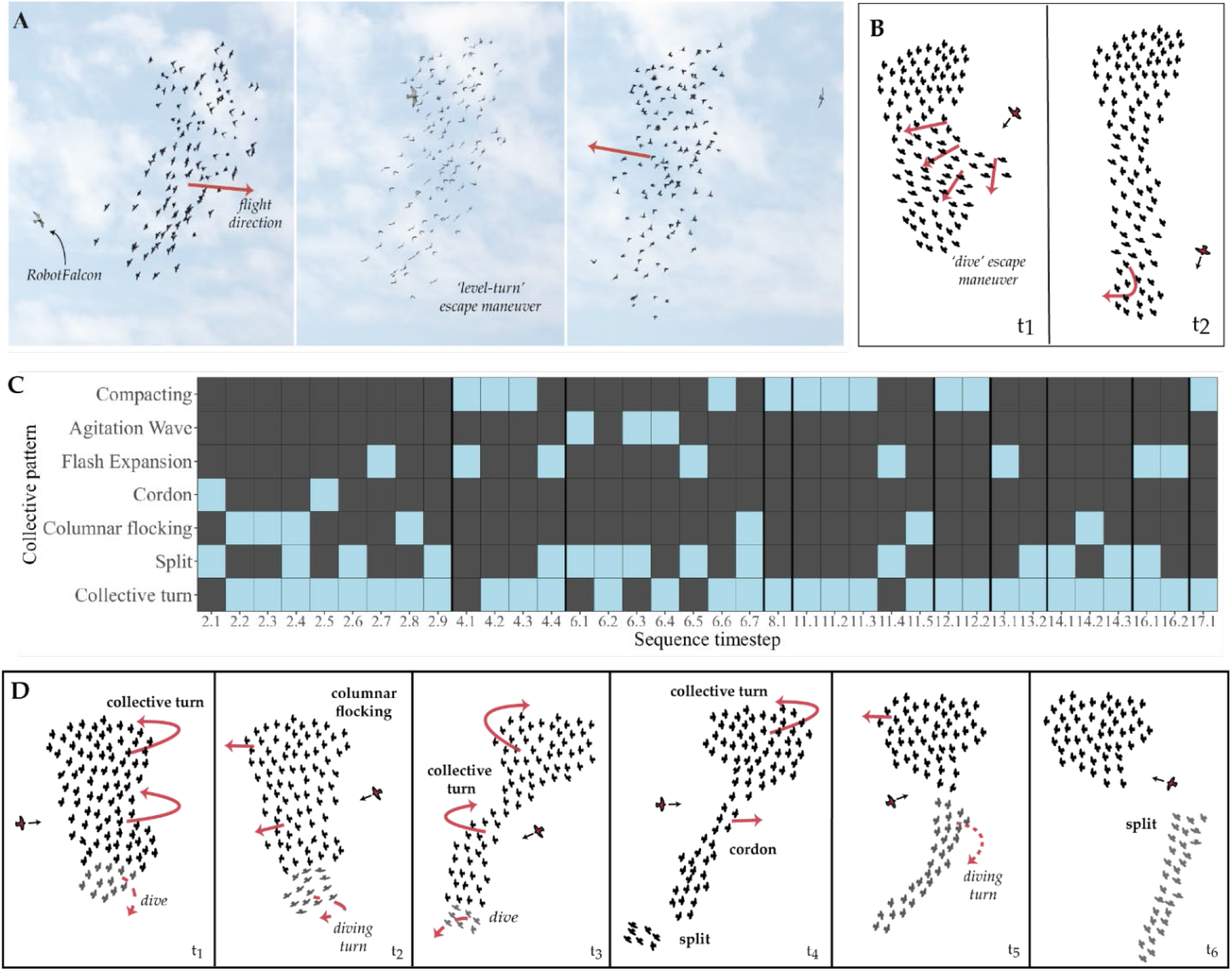
Empirical data of collective escape of starling flocks under attack by the RobotFalcon. **(A)** Screenshots of a flock performing a collective turn away from the RobotFalcon. Individuals perform a level-turn to evade the predator. The red arrows represent the flight direction of the flock. **(B)** Schematic representation of data of supplementary Movie S2B: individual starlings (small birds in black) dive away from the RobotFalcon (larger bird in red, with black arrow showing its heading). How much individuals dive seems to relate to their distance to the predator. A columnar shape of the flock emerges. **(C)** Co-occurrence of collective patterns in the empirical data. Every timestep (2.1 – 17.1) corresponds to a small time-window (∼1 second) in which certain patterns (light coloured blocks) appear in a given flock (defining a sequence). **(D)** Example of a sequence of different patterns of collective escape co-occurring (supplementary Movie S1), in which subgroups (parts of the flock) behaved differently. Each window (t_1_-t_6_) represents a snapshot during the sequence of collective escape. The initial configuration of the flock and relative position to the predator are given at t_1_. At t_2_, individuals that form the main part of the flock have performed a collective turn in the opposite direction (approx. 180°), while a small number of individuals at the bottom of the flock dived. At t_3_, individuals in the top part of the flock continue turning in the same direction, the ones in the middle part (closer to the predator) move slightly downwards (accelerating), while individuals at the bottom perform a diving turn. At t_4_, the main flock has performed a collective turn (approx. 180°). Due to the elongation of the flock, a cordon is formed. Individuals in the bottom part of the flock do not turn and continue to dive; as a result, they split from the main flock. At t_5_, individuals in the top part of the flock have performed another collective turn. Individuals in the middle and lower part continue their forward motion. At t_6_, the top part of the flock continues its forward motion while the bottom part performs a diving turn. As a result, the flock splits in two.

At the individual level, we observed starlings evading the predator by performing level-turns (reverting their flight direction without large changes in altitude, Fig. 1A), dives (changing their altitude away from the predator, Fig. 1B), and diving turns (turning while diving, Fig. 1D). All maneuvers are marked in the footage in supplementary Movie S2. We combined these observations with previous knowledge on the collective motion of starlings ^11,28,31^ to develop our new computational model of collective escape.

### StarEscape: an agent-based model of starling flocks

We built a spatially-explicit agent-based model based on self-organization to simulate the collective escape of starlings, named StarEscape. Modelling decisions as well as model parameterization and validation is based on the unique quantitative data of flocks in Rome ^10,11^.The model specifies two types of agents: ‘sturnoids’ that form flocks, and ‘predoids’ that chase them (inspired by the ‘boids’ of Reynolds 1987 ^24^). Agents move in a large (effectively infinite) 3-dimensional space. They are characterized by a position and a velocity vector in the global coordinate system, and their headings define the local coordinate system through which an agent senses the position and heading of its surrounding agents. Sturnoids have a set of 7 nearest neighbors (topological) ^9^ with which they interact through attraction, repulsion and alignment. Predoids follow the flock from behind for a given time period, after which they accelerate towards a single sturnoid, similar to the strategy of the RobotFalcon in the field. Agents `update’ their information about their surroundings asynchronously with a limited ‘reaction’ frequency. This reflects the behavior and cognition of birds in the wild: individuals in a flock neither react to changes in the position or heading of their neighbors instantaneously nor completely synchronously.

The individual behavior of sturnoids was split into 4 conceptual states: flocking, alarmed flocking, escape, and refraction. Transitions between the states are controlled by an individual-based and dynamic (non-homogeneous) Markov-chain. Overall, individuals coordinate with their neighbors when the predator is far away (flocking), and switch to alarmed flocking when the predator gets closer, during which they increase their reaction frequency, resembling increased coordination and vigilance expected when prey perceives predation threat ^44–47^. An individual performs probabilistically an escape maneuver depending on its perceived predation risk (based on its distance to the predoid, captured in its ‘stress’ level, see Methods) or on whether its neighbors are escaping (copying). After executing an escape, individuals remain in the alarmed flocking state but cannot perform another escape maneuver for a few seconds (refraction). Sturnoids perform two types of escape maneuvers that we observed in the starling flocks: level turns and dives. The selection of which maneuver to perform in the simulations presented here depends on a probability to turn (parameter *P*_*turn*_) or to dive (*P*_*dive*_ = 1 – *P*_*turn*_) (see also Supplementary text).

We analyzed simulated flocks of 20-5000 sturnoids (see supplementary Movie S3, S4, S6, and S7), with their assigned IDs referred to as SF (*Simulated Flock*). The flocks in simulations without a predoid (focusing on the collective motion of the flock, n = 18 independent simulated flocks) resembled flocks of starlings in their average speed (9.5 ± 0.32 m/s), nearest neighbor distance (NND = 0.86 ± 0.08 m, Fig. 2A) and stability of neighbors over a 3 seconds time window (Q_4_(3s) = 0.58 ± 0.06, Fig. 2B) ^10,28,48^. Other attributes of our simulated flocks such as scale free correlations in individual velocities (Fig.2C-2D), sharp flock borders, and variability of flock shape in our model is also similar to real flocks (Fig. 2E).

**Fig. 2.**
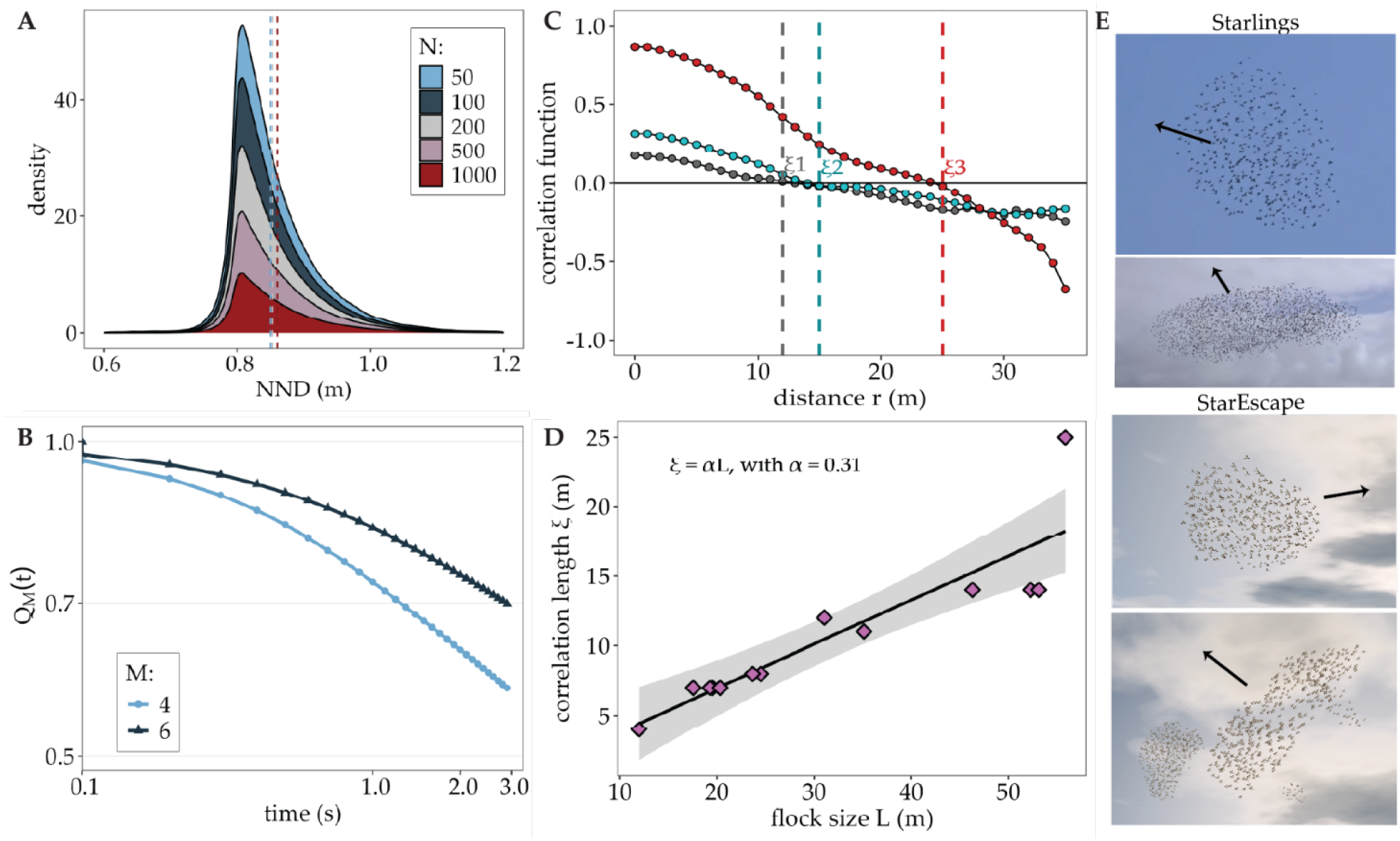
Flocking in StarEscape in the absence of a predator. **(A)** Distributions of nearest neighbor distance (NND) of simulated flocks (18) of different sizes (N = 50-5000). The vertical lines indicate the average NND for flocks of 50, 200, and 1000 individuals. **(B)** The average stability of neighbors (Q_M_, with M being the number of closest neighbors included in its calculation) over time in the simulated flocks (IDs *SF1-SF18*). **(C-D)** Scale-free correlations in simulations with 2000 (*SF19* and *SF20*, grey and blue points) and 5000 individuals (*SF21*, red points), calculated as in ^49^. In C, the vertical lines represent the correlation length ξ of each flock (ξ1, ξ2, ξ3 for flocks *SF19, SF20*, and *SF21*, respectively). In D, each point represents a simulated flock across 7 simulation runs (*SF19-25*), with the correlation length growing linearly with the size of the flock (Pearson’s correlation test: r = 0.9, p < 0.01). The timeseries of NND, Q_M,_ and velocity correlations are calculated by the model during a simulation. **(E)** Flocks of different sizes and shapes in starling flocks (top two) and the StarEscape model (bottom two). The arrows show the direction of motion of each flock.

### The emergence of collective patterns

Several patterns of collective escape emerged from sturnoids performing level-turning and diving maneuvers (Fig. 3). Note that the diving turn that we observed in the empirical data (Fig. 1D, t_2_ and t_5_) was not explicitly modelled, but diving turns emerged in our simulations when an individual reacted to the predoid with a turn without having levelled its pitch after a previous diving maneuver (see supplementary Fig. S4). Compacting emerged in our model from the flock becoming alarmed (with more frequent interactions among individuals) when the predoid was approaching (supplementary Fig. S5).

**Fig. 3.**
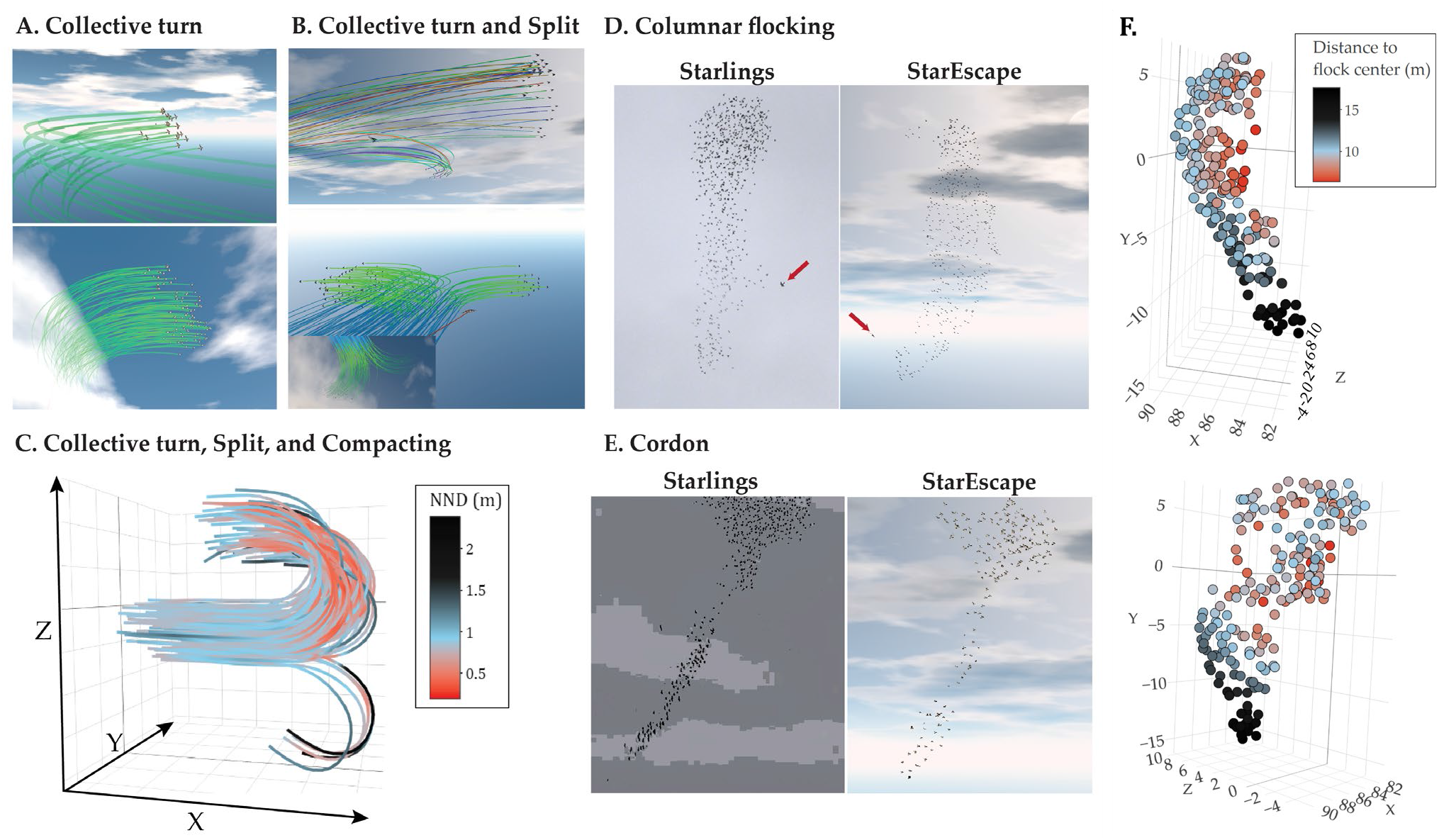
Patterns of collective escape in StarEscape. **(A)** Collective turns in flocks of 20 (top) and 200 (bottom) individuals, often co-occurring with **(B)** splits, and **(C)** compacting, emerging form individual level-turns. The trajectories and timeseries of nearest neighbor distance (NND) shown in (C) are exported by the model at the end of a simulation (N = 200, *Simulated Flock 26*). Altitude varies along the y-axis. From diving maneuvers, **(D)** columnar flocking and **(E)** the cordon emerge, similarly to what is observed in flocks of starlings pursued by the RobotFalcon. The red arrows in (D) indicate the position and orientation of the RobotFalcon and predoid (attacking a flock of 500 sturnoids). **(F)** The 3D coordinates of a flock (N = 200, *SF27*) during escape by diving. The color represents the distance of each individuals to the center of the flock as exported by the model.

In our simulations, collective turns of the whole flock propagated from a single or a few early responders with similar positions relative to the predator (Fig. 3A, supplementary Movie S3). Splits of sub-flocks emerged when a part of a flock was performing a collective turn and another part did not follow (Fig. 3B), i.e., when the level-turn did not propagate fast enough through the group (see also supplementary Fig.S6). A split emerged also when two early responders turned in opposite directions, being each followed by their own subset of neighbors (Fig. 3B-3C). This happened particularly when the predator approached from behind the middle of a flock, resulting in the two halves of the flock having different escape directions. Thus, sub-flock splits almost always co-occur with collective turns. Due to the copying mechanism, singletons rarely split off, and when they did, it was from the periphery of large flocks. During sharp escape maneuvers, blackening, compacting, and collective turn co-occurred (Fig. 3C). Blackening during collective turning may relate to both the relative orientation of the flock and the observer, and to an increase in group density.

Diving maneuvers usually led to columnar flocking and cordons (Fig. 3D-3F). Since the diving tendency of each individual increases when it is closer to the predator, columnar flocks emerged when the tendency of a flock member to dive resembled that of its neighbors. This is for instance the case in spherical flocks: the predoid is at a similar distance to a number of sturnoids at the back of the flock and the flock elongates vertically because these individuals that are closer to the predator reposition themselves downwards sequentially. If the predoid approaches the lower half of the flock, a dive may propagate only towards that lower part (supplementary Fig. S7). In this case, since flock members were still coordinating, there were a few sturnoids in the middle that were influenced by the position and heading of the diving individuals but did not dive themselves. As a result, these sturnoids, along with the top individuals of the diving subgroup, formed a ‘cordon’, connecting the two larger parts of the flock (Fig. 3E). If the predoid performed a fast attack at the bottom of the flock, a sub-flock split off, because a small number of individuals at the edge with higher predation risk rapidly dived. Collective turns while in columnar flocking, as observed in our empirical data (see Fig. 1A, schematic of Fig. 1D, and supplementary Movie S1), also emerged in StarEscape (supplementary Movie S4).

### Hysteresis

Expansion and compression of flocks in our model emerged through hysteresis: during a collective turn the flock often became denser which induced a subsequent decrease in density through repulsion forces among individuals right after the evasion (Fig. 4A-C). This emerges for a single parameter set of attraction, alignment, and avoidance (coordination) before and after the turn (Fig. 4B-C). The dynamics of this initial compacting may be linked to the differences in the initiation of turning across group members (supplementary Fig. S8). When some sturnoids start turning inwards, in relation to the flock’s centroid, they decrease their nearest neighbor distance, leading to more avoidance interactions after.

**Fig. 4.**
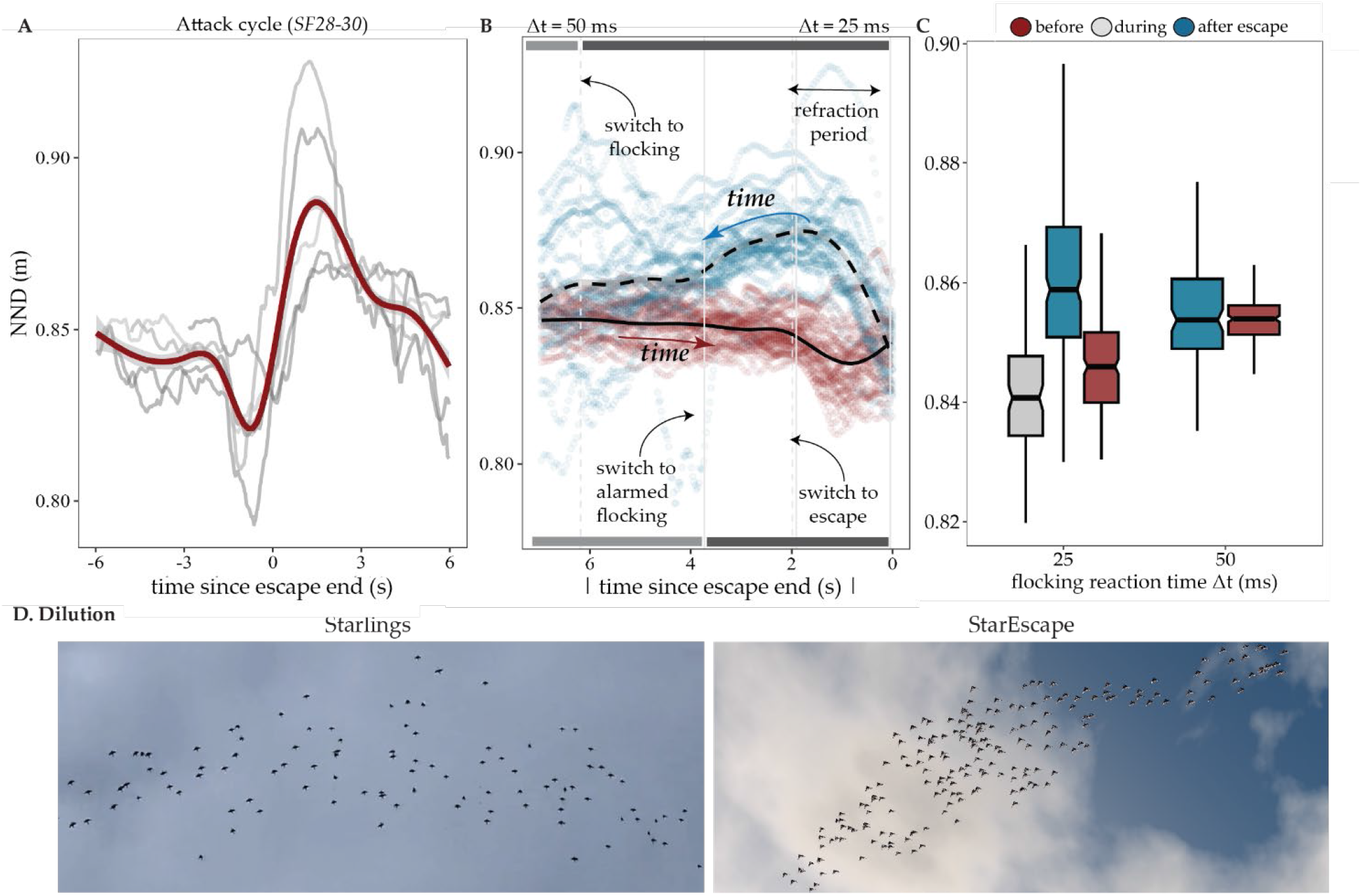
Hysteresis (collective memory) in collective escape. **(A)** Average nearest neighbor distance (NND) during an attack cycle of three simulated flocks (of 100 and 500 individuals, *SF28-30*). Time 0 is the end of a turning escape maneuver by all flock members. Grey lines show the time series of average NND during individual collective turns across the pursuit sequences. **(B)** The time series over absolute time relative to the end of an escape maneuver for 25 simulated flocks with 500 individuals (*SF29-53*). The points show the average NND of a single flock at each point in an escape cycle (the period in which the flock transitions from normal flocking to escaping the predoid and back). The solid line marks the changes in NND as flock members transition from regular to alarmed flocking (decreasing their reaction time from 50 to 20 ms), and then start performing a turning escape maneuver (‘before escape’ period). The dashed line shows the return to regular flocking (after escape), during which flock members have the same reaction frequency but higher NND as a result of the previous compacted state of the flock. The vertical lines indicate the average time that all group members have switched to a new state before (solid) and after (dashed) the end of the collective turn, respectively. Note that individuals do not switch between states synchronously but based on their own Markov-chains. The horizontal grey and black bars represent the reaction time of the agents before (bottom) and after (top) the end of the collective turn. **(C)** Average nearest neighbor distance in relation to the reaction time of agents during flocking (parameter Δt, in milliseconds) in simulated flocks (*SF28-96*). Despite the agents having the same rules of motion and interaction during alarmed flocking, their distance to their nearest neighbor differs before and after performing a collective turn. **(D)** The pattern of dilution emerges when individuals decrease their reaction frequency (switching from alarmed flocking to flocking) after the predator’s attack. Examples of dilution at the end of pursuits are also given in supplementary Movie S3 and S5.

The pattern of dilution in our model also emerged by hysteresis. It occurred as a side-effect of flock-members decreasing their reaction frequency when switching from alarmed flocking to regular flocking when the predoid retreated (supplementary Movie S5). Just from this decrease in coordination frequency, and with a denser flock pre-existing, the flock dilutes, taking a shape that differed from its shape at the initial condition before the predator’s attack (Fig. 4D). This matches the empirical observation that starling flocks dilute when they move away from the predator (Fig. 4D, supplementary Movie S5).

## Discussion

The first model of collective escape in swarms, namely fish schools, was published more than 20 years ago ^35^, but the complexity of the collective escape of starlings still makes their understanding particularly challenging. By combining empirical data and our new computational model, we show that the diversity of aerial displays of starlings under predation may arise from a combination of [1] a few individuals starting to perform an escape maneuver, [2] the speed with which this information transfers through the flock, [3] the local condition of each flock member (i.e., its position in the flock and its relative position to the predator) that determines its exact reaction, and [4] the relation between patterns (group shape and density) through time, i.e., hysteresis. Protocols for data collection in future studies should thus focus on those elements.

How the geometry of a group, that influences many effects of self-organization such as information propagation and hysteresis, affects its collective escape or decision-making has rarely been studied ^50,51^. An example of this is the columnar flocking we observed here. While the flock is columnar rather than spherical, the large differences in the relative position of individuals to the predator and the increased distance of all group members to the center of the flock (Fig. 3F) cause turning information to take longer to propagate from one edge of the flock to the other ^52^. As a consequence, when collective turns co-occur with columnar flocking, a cordon or a split may emerge, depending on the position and angular velocity of the escape initiator ^20^. The cordon tends to also lead to the split, since the two large parts of the flock experience different threat levels because of their different relative positions to the predator. It is interesting to note that the expected hunting strategy of predators of attacking the less dense part of a group (which during this formation is the cordon) to decrease confusion effects ^2,43,53^, also enhances splitting, which may further benefit the predator ^54^.

The specifics of escape maneuvers, information transfer, and local conditions we uncover here are valuable to also understand the emergence of other collective patterns of starlings, such as the agitation wave ^19,55^, the flash expansion ^8,18^ and the vacuole ^8^. The agitation wave can emerge from a zig-zag maneuver: individuals sequentially turning slightly to the left and then to the right (or vice-versa)^13^. Concerning the specifics of information transfer, previous work on agitation waves showed that more realistic waves (damping) emerge if individuals copy their neighbors’ maneuver with the same probability, but perform a less extreme maneuver (i.e., reduced banking) when they are further away from the predator ^14^. ‘Vertical’ compacting in starlings, a fast compression of the flock from the top that is not yet understood, may emerge from a similar mechanism: the reduced copying of a diving maneuver and the predator attacking the flock from the top. Understanding such mechanisms is also important for wave propagation in other systems (e.g., sulphur mollies, *Poecilia sulphuraria*, under attack by avian predators ^56,57^). The flash expansion can emerge from a fast turning maneuver away from the predator ^18,58^. According to both our empirical observations and previous findings in beetles ^58^, the flash expansion is a localized pattern, emerging as a direct response away from the position of the predator when the predator attacks at high speed and gets dangerously close to some individuals at the edge of a large flock (but see also ^34^ for a mathematical model on the emergence of flash expansions from multiplicative noise). The vacuole, a rare pattern seen only in very large flocks of starlings, is more common in fish schools (perhaps because of the lower maneuverability of their predators ^15,35,59^). It is characterized by some individuals being close to the predator without trying to escape. This may be due to the safety-in-numbers effect (risk dilution ^60^): the risk of an individual being caught is low in a large flock, so it may have a higher tendency to stay in the group and save energetic costs of maneuvering until the threat is too high. To better understand the emergence of more complex aerial displays, detailed information about escape reactions of prey individuals in relation to the predator’s position and speed ^61–67^, as well as their flock mates’ behavior ^27^, is necessary.

Our results further highlight that hysteresis may be crucial in the emergence of complex aerial displays. In collective motion of fish-like agents, a single parameter combination is shown to lead to both milling and polarized motion, depending on whether the behavior of the group before was ordered or disordered ^26^. We showed that a single parameter combination can lead to both a spherical compacted flock, and a wide, diluted flock (lower density), depending on whether the flock has been compacted or loose before. In self-assembling army ants, hysteresis has been identified when the gap that their bridge is covering is decreasing, with more individuals composing a bridge over the same distance than when the gap was increasing ^40^. In starlings, the effect of hysteresis we identified relates also to the fast sequence of temporary collective states of the flock during the whole sequence of a pursuit, while the flock transitions from noticing a predator, to escaping, and relaxing after the predator retreats. The temporal aspect of hysteresis (‘real-time’ collective memory) we identify here has to our best knowledge not been studied previously and may play a crucial role in the emergence of many spatiotemporal patterns (but see work on collective response to perturbations in swarms^68^), emphasizing the importance of considering long behavioral sequences in studies of collective escape and in the context of collective behavior over multiple time scales ^69^.

At the individual level, one aspect of behavior that is affected by the presence of the predator is the specifics of social interactions among prey, as known for fish schools ^44^. However, changes in the vigilance level of individuals, captured in agent-based models by the update or reaction frequency of each group member, has also been shown to lead to fast accelerations ^47^ (as identified in empirical data ^44^) and changes in the internal structure of the group (diffusion)^29^, without changes in the actual rules of social interactions. Our StarEscape simulations show that changes in the reaction frequency of group members may also underlie an emergent decrease in density, in line with what was expected to arise from the selfish herd hypothesis ^70,71^, but without an explicit increase in the tendency of individuals to move towards the center of their flock. This highlights the importance of considering such fine-scale effects of self-organization when interpreting collective phenomena.

Most of our current understanding of how patterns of collective escape emerge comes from computational models of fish schools ^17,35,59^. However, there are large differences between the locomotion type and evasive maneuvers of fish and birds, and in the internal structure of their swarms ^21,72^. Detailed investigation on how rules of interaction and escape ‘translate’ across systems is essential before we can establish universal mechanisms of collective escape. It is therefore unresolved whether similar collective escape patterns in fish ^16,35,59^ and starlings are generated by the same mechanism. Another limiting aspect of most existing models for studying starling flocks is their 2-dimensions. The importance of studying their flocks in 3-dimensions is apparent in our empirical data: by combining footage from the ground camera and the RobotFalcon we observe that the co-occurrence of compacting with collective turns may also be an effect of the observer’s orientation (supplementary Fig. S1)^13,16^. Our empirical observations suggest that co-occurrence of patterns emerges when the flocks are larger and the RobotFalcon pursuits the flock from closer and for longer. Which specific attributes are necessary for collective patterns to co-occur remains to be investigated.

StarEscape is a valuable tool to study the complex aerial displays of bird flocks in more detail, given also its user interface (supplementary Fig. S9), simulation speed, detailed output, and flexible structure facilitating future development. Further model functionalities not used in the analysis presented here, such as the deterministic or dynamic selection of escape maneuver, are mentioned in our Supporting Information Text. In terms of collective motion, StarEscape can be extended to investigate other characteristic of starling flocks for which underlying mechanisms are not well understood, such as the emergence of high density borders. Concerning future work on collective escape, a limiting aspect is the difficulty to investigate the emergence of different collective patterns across the parameter space of our model. To overcome this constraint, automated identification of the different patterns should be developed, i.e., a quantitative definition of each type of collective escape, similar to distance-based thresholds used for splitting in pigeons ^20^ and proxy metrics used to study patterns of collective motion such as turning ^29^ and milling ^30^. The existing monitoring of escape propagation networks, predator-prey details (e.g., relative position of each sturnoid to the predoid), splitting, collective turning, velocity correlations, and flock shape (based on object-oriented box approach) support future endeavors to quantitatively investigate the mechanisms by which flocks transition between collective states and thus quantify self-organized dynamics.

Apart from this process-oriented investigation, a function-oriented analysis in StarEscape can show the adaptiveness of different patterns, how they affect the catching success of the predator ^2,17^, or their dependence to individual heterogeneity in terms of escape reaction ^73^ (since selection pressures are expected to act on such response parameters). StarEscape can further be adjusted to other bird species, such as dunlins ^43^, since the coloration and shape of agents in our model can be changed (see supplementary Movie S6), allowing across species comparisons ^16,72^. For instance, the ‘rippling’ effect that dunlin flocks demonstrate ^43^ may be the result of orientation waves similar to the ones of starlings ^13,43^. Overall, a comparative approach over several patterns of collective escape in 3-dimensional groups can improve our understanding of starling murmurations, complex aerial displays in general, and the evolution of such extreme collective phenomena.

## Methods

### Experimental design and empirical data analysis

We analyzed video footage of 19 flocks of European starlings (*Sturnus vulgaris*) pursued by our RobotFalcon ^39,41^. The dataset was collected along with data on other bird species (corvids, gulls and lapwings; by Storms et al. (2022)^39^) between February and November 2019 at agricultural sites in the north of the Netherlands (52°59’N-5°27’E). The experimental work was carried out in agreement with the rules of the University of Groningen Animal Welfare Committee. We have complied with all relevant ethical regulations. The RobotFalcon approached flocks from a distance while the flock was on the ground (stationary) and continued to an aerial pursuit when the flock took off, until it left the site. Flocks that were already airborne were not approached. The RobotFalcon is a remotely controlled aerial predator that resembles a peregrine falcon (*Falco peregrinus*) in appearance and size. It is steered by a trained pilot, aiming to imitate the hunting behavior of the real falcon. The pilot has a first-person view of the flock through a camera mounted on the back of the RobotFalcon. To follow the flocks during their full collective escape sequences, the ground camera has to move, prohibiting the use of methods based on stereophotography with three view-points to retrieve individual tracks for short time periods^10,74^.

In the footage retrieved from the ground camera, we identified the patterns of collective escape according to previous classifications ^7,8^, with the exception of the split. We identified a split when part of the flock formed a new cohesive flock, without taking into account splits of one or a few individuals (as a result of a flash expansion for instance, as done in previous work of small groups of 10-30 individuals)^20,33^. We further recorded whether in a single flock different patterns of collective behavior were exhibited simultaneously. In detail, we analyzed videos (N = 19) of approximately 15 minutes of pursuit by the RobotFalcon using the Adobe Premier Pro software and using the classification system of previous work on starling flocks under attack for identifying collective patterns^8,41^. In detail, we factorized each video into a sequence of steps during which a new collective state is observed (a specific pattern appears or disappears, e.g. collective turns last for approximately 1 second). Periods in between patterns of collective escape, during which the flock remained stable in shape, color, and direction of movement, were ignored. From all the derived timeseries of collective events, we kept only the flocks in which two collective patterns co-occurred in at least one step (N = 10). Footage from the RobotFalcon camera was used to compare the observed patterns from a different angle (to investigate viewer’s perspective illusions, e.g., Moiré effects^69^), when possible (supplementary Fig. S1). For each event of collective escape, we also examined the motion of individual starlings that were close to the border of their flock, particularly focusing on changes in their heading and altitude. We selected few visible individuals close to the point of attack of the RobotFalcon and tracked their motion in the two visible dimensions. We used our observations to inspire the development and guide the calibration of an agent-based model.

### A computational model of collective escape

We built an agent-based model to simulate the collective escape of starlings (StarEscape)^82^. The model is based on the principles of self-organization with two types of agents moving in an open 3-dimensional space: ‘sturnoids’ that coordinate with each other and ‘predoids’ that pursue and attack the sturnoid flocks. We present our model below, using elements of the ODD protocol ^75^, but see our Supplementary text for a complete model description. The model is written in C++, using OpenGL ^76^ for the real-time visualization of the simulation, and DearImGui ^77^ for the graphic user interface (GUI, supplementary Fig. S9).

### Model overview

All agents are defined by a position and a velocity vector in the global coordinate system. The heading of each agent defines its local coordinate system (Fig. 5A) through which an agent senses the position and heading of its interacting agents. Sturnoids have a set of nearest neighbors with which they interact (set ℕ) and predoids have a flock or a sturnoid as a target. According to empirical findings on starling flocks, we model topological interactions between individuals ^9^. Each sturnoid interacts with its 7 closest neighbors (*n*_*topo*_) that are within its field of view (*θ*_*FoV*_) according to the rules of alignment, attraction and avoidance (flocking sub-model^75^).

**Fig. 5.**
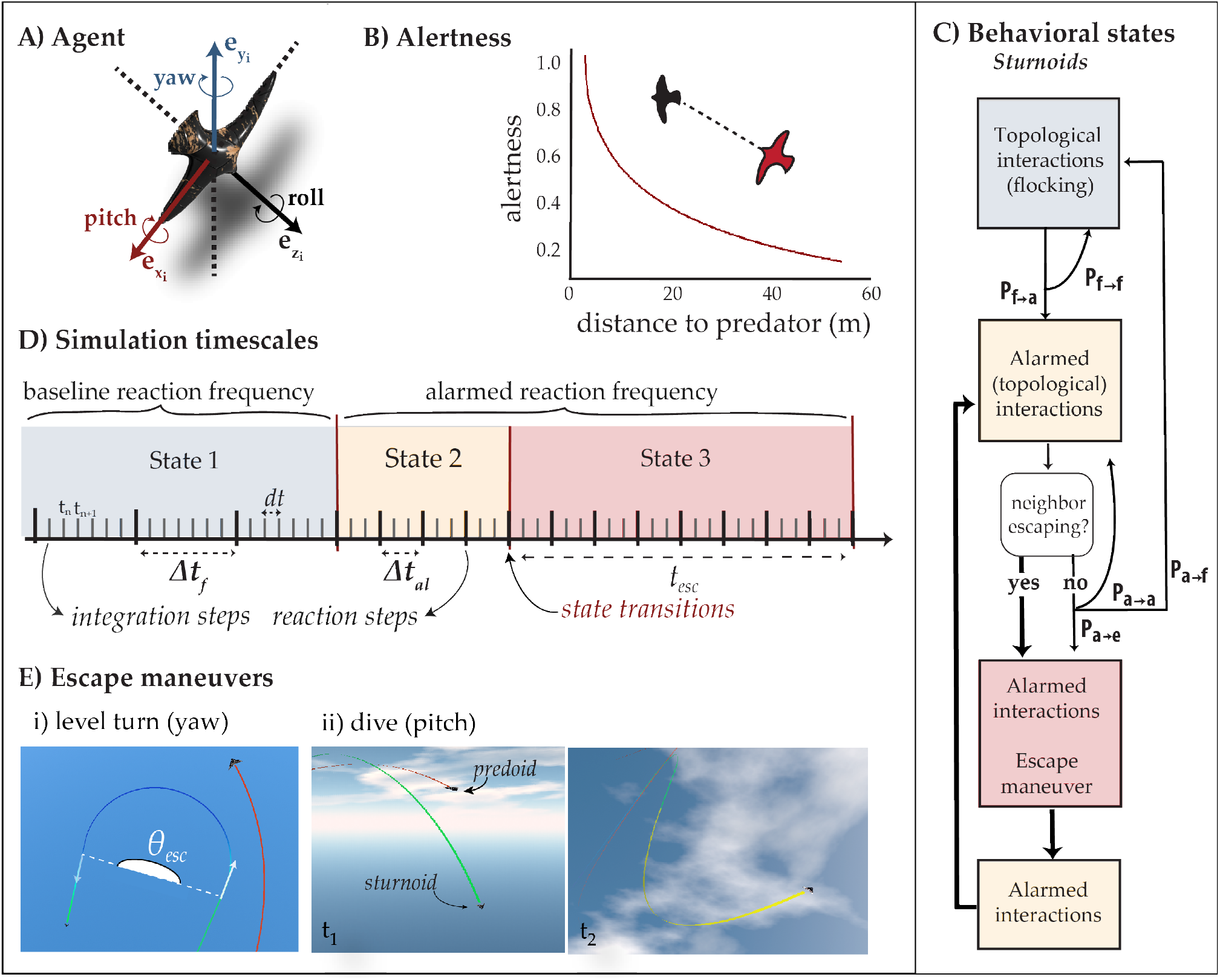
The StarEscape model. **(A)** The local coordinate system of an agent in the 3-dimensional space. The vectors e_x_, e_z_, and e_y_ represent the sideward, forward, and upward directions of the agent, respectively. The agent yaws and rolls to turn, and pitches to dive or climb. **(B)** Alertness increase. The alertness of each individual accumulates at each update timestep as a function of its distance to the predator (see also supplementary Fig. S12 for example data). **(C)** The transition mechanism between behavioral states in sturnoids. The transition probabilities (P_x->x_) represent the non-homogeneous Markov-chain of each individual and thicker arrows represent deterministic transitions. In detail, sturnoids may transition from their regular coordination (with update frequency *1/Δt*_*f*_) to an alarmed state if their alertness increases. During alarmed coordination, their coordination frequency (*1/Δt*_*al*_) is increased and a sturnoid may perform an escape maneuver based on its probability to escape or copy the escape of its neighbors (transitioning to an escape state). Each escape has a predefined duration (*t*_*esc*_), after which agents transition back to alarmed coordination without the option to perform an escape maneuver (refractory state). After this, they regain the probability to react to the predator and the cycle starts again. If alertness decreases, individuals transition back to regular coordination. The exact values of the Markov chain depend on each individual’s alertness. We derive them by interpolating through 3 parameterized transition matrices for low, intermediate, and high alertness. The chain of the predoid is given in supplementary Fig. S13. **(D)** The timescales of StarEscape. At every integration step (t_n_) individuals update their position and heading according to their steering force (with frequency *1/dt*). At every reaction step, individuals recalculate their steering force, representing the cognitive collection and processing of information from their environment (with frequency *1/Δt*). At these time steps, each individual also decides whether to stay in its current state or switch, depending on its Markov-chain. **(E)** Screenshots of the escape maneuvers of individual sturnoids in a simulation. Sturnoids escape the predator by two evasive motions, (i) a level turn (with angular change *θ*_*esc*_) or (ii) a dive (t_1_, after which they return to level flight, t_2_). The blue and green tracks show the sturnoids’ trajectories during escape, the red tracks show the predoid’s trajectories, and the yellow track the refractory period and pitch recovery after the diving maneuver.

Apart from their neighbors, sturnoids sense the position of the closest predoid and their ‘alertness’ (or ‘stress’) increases as they get close to it (Fig. 5B). Sturnoids react to the predoid with an evasive maneuver based on a probability that depends on their alertness (similar to decision-making processes in models of evidence accumulation in social systems ^78^). The neighbors (*n*^*topo*^_*cop*y_) of an escaping sturnoid copy its maneuver. A predoid first chooses a flock and follows it from a given point relative to its center (based on the parameters *β*_*at*_ and *d*_*at*_). From there, it picks its closest sturnoid as its target and attacks it. The attack has a specific duration (*t*_*hunt*_) after which the predator gets re-positioned far away from the flock. Since we focus on the emergence of collective escape patterns, we do not model ‘catches’ of sturnoids by the predator.

### Behavioral states

The individual-level rules of motion and interaction in our model depend on the ‘state’ of each agent. For sturnoids, we define 3 behavioral states: a coordination state, an alarmed-coordination state, and an escaping state (Fig. 5C). An individual-based non-homogeneous Markov Chain controls the transition from one state to another (see example in supplementary Fig. S10). The transition probabilities depend on the alertness of each individual at each timestep, which depends on its distance to the predator. When a sturnoid is close to the predator, its alertness increases every time it updates its information about the predator’s position, according to a given function (see section ‘Escape’ and Fig. 5B), and decays with a constant rate (*r*_*decay*_). The increase and decay of alertness is balanced for alertness to always increase when the predator is close and decrease back to 0 within a few seconds after the predator retreats (supplementary Fig. S11). From this mechanism, individuals switch to alarmed coordination (higher reaction frequency) when the predator starts pursuing the flock, spontaneously escape (depending on alertness) when it gets close, and return to their basic coordination when the predator stops its pursuit.

During coordination, group members align with, are attracted to, and avoid each other (for details see section ‘Coordination’). Sturnoids are also attracted to their roost and to a given altitude (see section ‘Physics of motion’). During alarmed coordination, individuals coordinate with each other more often (higher reaction frequency, *Δt*_*al*_), according to empirical evidence that prey individuals have faster reactions under predation (more frequent interactions with neighbors)^44,47^. When alarmed, individuals may switch to an escape state. This switch is either an individual ‘decision’ (based on the probability of their Markov chain) or induced by an escaping neighbor; individuals copy the escape reactions of their interacting neighbors. During escape, sturnoids perform an evasive maneuver while coordinating, with the relative influence of each term (e.g., alignment, attraction, turn away) being controlled by their relative weights. After escaping, individuals return to an alarmed-coordination state without the possibility to escape for a specific period of time (conceptualized as a refractory period, *t*_*ref*_). This chain of behavioral states and its transitions are given in Fig. 5C. More details about the transition probabilities are given in the Supplementary text.

The predoid in our simulation has 3 behavioral states. First, it perceives the largest flock (in case a split has occurred) and positions behind it at a given distance (*d*_*at*_) and bearing angle (on the horizontal and vertical plane, *β*_*at*_) from the flock’s center and velocity, adopting its heading (preparation state). Second, it follows the flock (‘shadowing’) or locks into its closest sturnoid and moves towards it, with a speed that scales from the target’s speed (*u*_*scale*_, e.g., 1.3 times faster than the sturnoid; attack state). Third, after the attack duration has passed (*t*_*hunt*_), the predator repositions away from the flock for a given time period (retreat state, *t*_*retreat*_). The cycle then repeats, and a new attack takes place. Thus, all the states of the predoid (see supplementary Fig. S13) have predefined durations, like the refractory period of sturnoids and opposite to the rest of the sturnoids’ states that spontaneously alternate.

We further describe the sub-modules of each state below. To better understand their specifics, we first explain the behavioral timescales in our model and how individual motion is controlled.

### Timing

In the model, time is measured in seconds according to a parameter that scales a time step of the model to the real units of time (*dt*). We use two timescales, the ‘integration step’ and the ‘reaction step’. A single time step (*dt* seconds) is an ‘integration’ step. Since individuals cannot instantaneously receive, process, and react to information about their environment, we define as ‘reaction step’ the frequency with which individuals update their information about their environment (neighbors and predator). Every reaction step consists of several integration steps, during which agents integrate the information they collected in the preceding reaction step by adjusting their position and heading (Fig. 5D). Since grouping individuals in nature are not perfectly synchronized, individuals in the model update their information asynchronously (at different integration steps) but with the same frequency. During a reaction step, individuals may also switch state as described above.

### ‘Physics’ of motion

Individuals are steered in the global space by a pseudo-force (referred to as ‘steering force’): a 3-dimensional vector that pushes individuals to their future position and heading. The steering force has 4 components: 1. a coordination force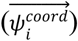, 2. an escape force (for sturnoids;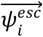) and a pursuit force (for predoids;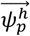), 3. a control force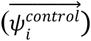, and 4) a drag force 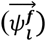. Large values in the forward direction (z) of the steering force will make individuals accelerate, towards the side (x) will make them turn, and in the vertical direction (y) will make them change altitude (dive or climb). An update force is calculated at each reaction step as the sum of all components included at a given state:

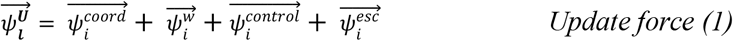

with the control and escape vectors being 0 during escape and flocking states, respectively. At each integration step, the total steering vector is calculated by:

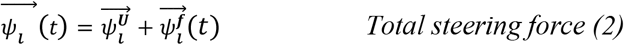

This steering force is then used to calculate the acceleration of each agent according to Newton’s second law of motion and calculate at each integration step the agent’s new position and heading (using the midpoint method, see Supplementary text). Details about the drag and control force are given in the Supporting Information Text.

### Coordination

The specifics of coordination affect the motion of a sturnoid continuously. Attraction pulls each individual towards the center of its neighborhood (*n*^*topo*^_*coh*_). The strength of this force depends on the individual’s distance to that center: the further away a sturnoid is from its neighbors’ centroid, the stronger the force is. Alignment turns an individual towards the average heading of its neighbors (*n*^*topo*^_*ali*_). Avoidance makes the individual turn away from the position of its closest neighbor, if the neighbor is within a range of minimum separation (*d*_*s*_). We parameterize each sturnoid to avoid only its closest neighbor (instead of their *n*^*topo*^ closest neighbors) for the diffusion of flock members in our simulations to resemble more empirical data of starlings^28^. Noise to each individual’s motion is also added through a pseudo-force perpendicular to its heading with magnitude sampled from a uniform distribution with bounds *-w*_*n*_ and *w*_*n*_ (weighting factor for random noise). Details on the calculation of each component are given in the Supplementary text.

### Escape

At every reaction step, the alertness of an individual accumulates according to the function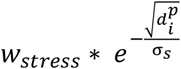 (with 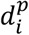being the distance between the focal individual i and the predator, *σ*_*s*_ a scaling parameter, and *w*_*stress*_ the weighting factor for stress accumulation), and at every integration step it decays (with rate *r*_*decay*_). The higher an individual’s alertness is, the higher its probability of performing an escape maneuver. A parameterized set of probabilities assigned to each maneuver (*P*_*x*_) controls which evasive action the individual will perform (but see also *Further functionality* subsection in Supplementary text). We investigate two maneuvers in our model: a level turn and a dive (Fig. 5E). During turns, individuals make a turn of *θ*_*esc*_ degrees (here ≃180^°^) away from the predator (depending on their own heading relative to the predator’s heading; see Supplementary text). During dives, individuals pitch downwards depending on their own distance to the predator (the closer the predator the steeper the dive). This distance dependency is controlled by a smootherstep function ^79^:

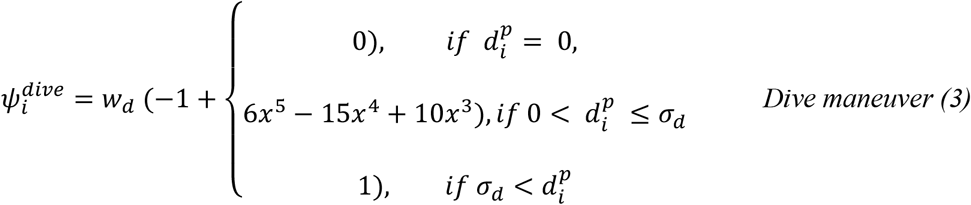

where,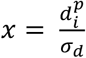, *w*_*d*_ is the weighting factor for diving, and *σ*_*d*_ the smoothing parameter that denotes the minimum distance from the predator for a sturnoid to dive. When an individual exceeds a maximum dive distance (*d*_*d*_), it starts pitching upwards to get to its preferred altitude. Note that during copying, an individual copies the type of escape maneuver from any of its interacting neighbors (turn or dive), but bases the specifics of the maneuver on its own local conditions (relative position and heading to the predator). The topological range for copying (*n*^*topo*^_copy_) is set to the same value as for coordination. In the event that an agent has more than 1 neighbor performing a different maneuver, it copies the maneuver of its closest escaping neighbor.

### Simulation design, data collection and analysis

Our simulations include a single predoid and many sturnoids (N <=5000). Sturnoids are initialized in a flocking formation: with a small heading deviation around a given global heading and positioned within a radius around a given point in space. This resembles the starting conditions of the starling flocks in our empirical experiment. The predoid is initialized a few seconds after the flock and immediately enters its preparation state (positioned behind the flock with the same heading as its target). All parameter values (supplementary Table S1) are given to the compiled model through a configuration file (.*json*) at run time (for the selection of the default values see our Supporting Information Text). We focus our analysis on which patterns emerge across simulations. A camera that follows the flock (with a certain flock member as its center) shows the simulation in real time. The user can move the orientation of the camera to look at the flock from a different perspective (see supplementary Fig. S9 and Movie S7).

First, we collected screenshots and recordings of the emerging patterns of collective escape across simulations. We compare these observations with the empirical data of starlings’ collective escape (video recordings). Second, we analyzed a number of measurements computed by the model in real time and exported at the end of each simulation (for details see our Supporting Information Text). The available quantitative data of starling flocks are very limited, constraining comparisons of our model across many characteristics of starling flocks. However, to validate the collective motion in our model, we run simulations without the predator, and compared the average speed, nearest neighbor distance, stability of neighbors (based on the 4 and 6 closest ones, measured as in ^28,29,48^) and velocity correlations (as measured in ^49,80^) of our simulated flocks (N = 50-5000) with previously published empirical results from starlings in Rome ^10,11,48^. This pattern-oriented approach ^22^ has been shown to successfully accomplish the resemblance of simulated flocks to the target species in comparisons over a dozen metrics of collective motion ^32,81^. For our hysteresis analysis, we further investigated the dynamics of group density during collective escape, using trajectories from 75 simulations with 90 second pursuits (according to the average pursuit duration of the RobotFalcon) on flocks of 100-500 individuals.

### Statistics and Reproducibility

All observed flocks of starlings (F) and simulated flocks (SF) were assigned a unique ID. The configuration files used to produce the simulated data are provided along the simulated trajectories and data of each flock (n = 96)^83^. Further data analysis, statistics, and visualization were performed with custom code in R^84^.

## Acknowledgments

The authors would like to thank the group of self-organization of social systems at the University of Groningen for their support and comments at various stages of this project, as well as Robert Musters for building and piloting the RobotFalcon, the students Ronja Hulst, Sorscha Passmore, and Deborah Salleh for contributing to the collection of the empirical data, and Heleen Jansen for contributing to a pilot analysis of the video footage. This work has been financed by the Netherlands Organisation for Scientific Research (NWO - https://www.nwo.nl), the Open Technology Program (OTP) “Preventing bird strikes: Developing RoboFalcons to deter bird flocks” (project number 14723) awarded to C.K.H. C.C. and M.P. have also been supported by the Italian project PRIN 2020 “Collective and individual responses of avian flocks to robotic predators” (2020H5JWBH) in collaboration with C.K.H.

## Author Contributions

MP, HH, and CKH conceptualized the study. RS collected the empirical data, supervised by CKH, CC and SV. MP and RS analyzed the empirical data with input from CC, SV and CKH. MP and HH developed the computational model. MP designed the computational experiments, ran the simulations, analyzed the simulated data, created all visualizations, and wrote the original manuscript. All authors provided feedback and revised the manuscript.

## Data and Code Availability

The empirical and simulated data that support the findings of this study are available in Zenodo with the identifier(s) [DOI to be added upon publication]^83^. The computational model StarEscape developed in this study and the custom code used for the data analysis are available on the GitHub repositories https://github.com/marinapapaStarEscape-model and https://github.com/marinapapa/CollectiveEscapeStarlings, respectively, and the versions used for this study are archived in Zenodo with the identifiers [DOI to be added upon publication]^82^ and [DOI to be added upon publication]^84^, respectively.

