## Supplementary Material for "Towards a mechanistic understanding of collective escape in starlings"

**This PDF file includes:**

Supplementary Text
Figs. S1 to S13
Tables S1
Movies S1 to S7 (link and captions)

**Other Supplementary Materials for this manuscript include the following:**
Movies S1 to S7

### Supplementary Text

#### The StarEscape model

We present our 3-dimensional agent-based model, StarEscape, following the terminology of the ODD (Overview, Design concepts, Detail) protocol [*1*]. The model is built in C++ 17, using OpenGL *[2]* for visualizing the simulations and DearImGui *[3]* for user interface. Simulations can be run 'headless' without the visualization to increase speed.

The model consists of sturnoids and predoids; two types of agents imitating the behavior of an individual starling (*Sturnus vulgaris*) and of an attacking predator respectively. All agents have a position and a velocity vector defining their placing and heading in the simulation space, a three-dimensional large cubic space, representing the aerial space. Space is measured in meters. Each simulation has multiple sturnoids and one predoid. Each time step of the model represents *dt* seconds of real time (see Main Text).

Coordination

Each sturnoid can sense the position and direction of a specific number of neighbors within its field of view. Individuals positioned in the blind angle at the back of each sturnoid are not sensed by the focal agent. The model is based on the rules of topological (closest $n_{topo}$ individuals) attraction, alignment and avoidance, adjusted to the species-specific behavior of starlings. We model centroid-attraction (a force towards the center of mass of each topological neighborhood) scaled by the distance of the focal individual to this centroid to ensure sharp borders that are a characteristic of starling flocks (see also [*4*, 5]). Specifically, the vector for attraction is calculated as:

$$\overrightarrow{v_i^c} = \frac{1}{n_{topo}} \sum_{j \in N_i(t)} (\overrightarrow{r_j} - \overrightarrow{r_i}) \quad (1)$$

where $\vec{r}_j$ is the position vector of a neighbor j, and $N_i(t)$ the set of topological neighbours at time *t*. The weight of the steering force is then adjusted according to a smootherstep function:

$$w'_{ci} = w_c * \begin{cases} 0, & if\ x < \sigma_{coh}^{l} \\ 6x^5 - 15x^4 + 10x^3, & if\ \sigma_{coh}^{l} \le x \le \sigma_{coh}^{h} \\ 1, & if\ x > \sigma_{coh}^{h} \end{cases} \quad (2)$$

with $x = |\overrightarrow{v_i^c}|$, the distance between the focal individual and the centroid of its topological neighbors. The total attraction component of the steering force is then calculated from:

$$\overrightarrow{\psi_i^c} = w'_{ci} * \frac{\overrightarrow{v_i^c}}{|\overrightarrow{v_i^c}|} \quad (3)$$

Alignment is based on the average heading ($\hat{h}$) of each topological neighborhood:

$$\overrightarrow{\psi_i^a} = w_a \ * norm( \sum_{j \,\epsilon\, N_i(t)} \widehat{h_j} ) \qquad (4)$$

with *norm* indicating a vector normalization function $norm(\vec{x}) = \frac{\vec{x}}{|\vec{x}|}$ .Avoidance is based on the position of only the nearest neighbor (parameter $n^{topo}{}_{\underline{sep}}$), if placed within a minimum separation distance:

$$\overrightarrow{v_i^s} = (\overrightarrow{r_i} - \overrightarrow{r_{nn}})$$

and

$$\overrightarrow{\psi_i^s} = w_s \begin{cases} 0, & if \ |\overrightarrow{v_i^s}| > d_{min}^s \\ norm(\overrightarrow{v_i^s}), & if \ |\overrightarrow{v_i^s}| \leq d_{min}^s \end{cases} \qquad (5)$$

where $\vec{r}_{nn}$ is the position of the nearest neighbor from $N_i(t)$, $d_{min}^s$ the minimum separation distance between two individuals, and $w_s$ the weight for avoidance.

The total coordination steering force is composed as:

$$\overrightarrow{\psi_i^{coord}} = \overrightarrow{\psi_i^c} + \overrightarrow{\psi_i^a} + \overrightarrow{\psi_i^s} \qquad (6)$$

*Emergence*

Based on the coordination among sturnoids, the agents group together forming flocks. A sturnoid belongs to a flock with those individuals with whom it shares a neighbor within 10 meters. Multiple flocks can be formed during a simulation, by some individuals splitting from the group. Each flock is assigned a unique ID. By the sturnoids maneuvering away from the predoid, different patterns of collective escape emerge.

*Predator-prey interactions*

Predoids sense the position and speed of the sturnoids. A predoid can also distinguish which flock in the simulation is larger and pick a member of this flock to follow as its target. Sturnoids sense the position and heading of their closest predoid.

Sturnoids interact with the predoid by performing one of two available escape maneuvers: a level-turn or a dive. The turning angle and duration of the level turn are sampled from two gamma distributions independently. The force needed for the sturnoid to perform this turn across multiple timesteps depends on its speed at the beginning of the maneuver and the calculated radius of the turn according to:

$$\gamma_\iota = \frac{u_i \, T_E}{\theta_e}$$

and

$$\overrightarrow{\psi_\iota^{esc}} = \frac{{u_i}^2 \, m}{\gamma_\iota} \, \widehat{\psi_\iota^p} \qquad (7)$$

where $T_e$ and $\theta_e$ are the sampled duration and angle of the maneuver respectively, $u_i$ the speed of the focal individual, and $\widehat{\psi_\iota^p}$ the direction away from the predator. The level turn is performed in the x-z plane of their reference frame (note not the global reference frame); the sturnoids don't change their pitch.

The dive is performed in the z-axis, away from the predoid's position. The closer the predoid the deeper the sturnoid's dive (see main text for motion equation). A sturnoid performs a maneuver depending on a probability that increases with decreasing distance to its position. The exact maneuver is selected based on a parameterized probability ($P_{turn}$, $P_{dive}$). During an escape maneuver, the sturnoid still coordinates with its neighbors.

The predoid interacts with the sturnoids by following them from a given distance and relative position (`pursuit' or 'shadowing', controlled by parameters $d_{at}$ and $\beta_{at}$) with speed scaling from the flock's speed ($u_{scale}$). The pursuit lasts for a given duration ($t_{hunt}$), after which the predoid is repositioned far away from the flock for a given duration ($t_{retreat}$), before being repositioned back to start a new pursuit (see also supplementary Fig. S8B). The pursuit force of the predoid is calculated through:

$$\overrightarrow{\psi_p^h} = w_h * norm\left(\left(\overrightarrow{r_z} + d_{at}\boldsymbol{R}\widehat{h_z}\right) - \overrightarrow{r_p}\right) \qquad (8)$$

where $\boldsymbol{R}$ is a rotation matrix, $\widehat{h_z}$ the heading of the targeted nearest prey from the largest flock, and $\overrightarrow{r_p}$ and $\overrightarrow{r_z}$ the position vectors of the predator and the nearest prey respectively.

The predoid cannot `catch' a sturnoid, and an agent cannot die.

<u>Stochasticity</u>

At each update step, there is a random error added to the motion of each agent. It is represented by a 'wiggling' force:

$$\overrightarrow{\psi_\iota^w} = \varepsilon_h \widehat{h_\iota^\perp} \qquad (9)$$

with $\varepsilon$ a noise scalar sampled by a uniform distribution based on the parameter $\underline{w_n}$, and $\widehat{h_\iota^\perp}$ the unit vector perpendicular to the heading vector of the focal agent.

The decision of a sturnoid to perform an escape maneuver and the selection between level-turn and a dive are stochastically chosen. The probability of displaying a maneuver increases with decreasing distance to the predoid (supplementary Fig. S11). The total turning angle and duration of a level-turn are each sampled from a gamma distribution.

*Physics of motion*

Apart from the coordination and escape (or pursuit) steering forces (as described in the main text), a control and a drag force are modelled to imitate the flying of birds in nature. The control force is composed by, first, a level attraction force ($\psi_i^{pitch}$) that represents the inability of birds to climb sharply (especially when they are part of a flock). The level force constrains the pitch of an individual to a given maximum ($p_{alt}$). We also model an 'altitude attraction': individuals have a preferred altitude ($y_{alt}$) towards which they tend to return according to a smootherstep function *[6]*:

$$\psi_i^{alt} = w_{alt}\, \xi(-1 + 2 \times \begin{cases} 0), & if -\sigma_{alt} > d_i^{alt} \\ 6x^5 - 15x^4 + 10x^3), & if -\sigma_{alt} \leq d_i^{alt} \leq \sigma_{alt} \\ 1), & if\ \sigma^{alt} < d_i^{alt} \end{cases} \qquad (10)$$

where $x = \frac{d_i^{alt}}{\sigma_{alt}}$, $\sigma_{alt}$ is the smoothing parameter around the preferred altitude, $w_{alt}$ the weighting factor of altitude attraction, $\xi = -sin\,(p_{alt})$, with $p_{alt}$ the maximum pitch for altitude attraction (in rads), and $d^{alt}{}_i$ the absolute deviation of the altitude of individual $i$ from the preferred altitude. To keep our flocks within our simulation space, we define a 'roost' inspired by real flocks *[5, 7, 8]*. It is a point ($r_r$) on the plane (x-z) towards which sturnoids are attracted if they get far away from it, with weighting factor $w_r$ and according to a smootherstep function based on their distance to the border of the roost:

$$\overrightarrow{v_i^{roost}} = (\overrightarrow{r_r} - \overrightarrow{r_i})$$

and

$$\overrightarrow{\psi_i^{roost}} = w_r \begin{cases} 0, & if\ x < \theta_r{}^2 \\ 6x^5 - 15x^4 + 10x^3, & if\ \theta_r{}^2 < x \leq \zeta \\ 1), & if\ \zeta < x \end{cases} \qquad (11)$$

with $x = \left|\overrightarrow{v_i^{roost}}\right|^2$ , ζ being a hard-corded large constant (500,000), $\theta_r$ the radius of the roost (around point $r_r$). The total control force is then calculated as:

$$\psi_i^{control} = \psi_i^{alt} + \psi_i^{pitch} + \psi_i^{roost} \tag{12}$$

It is important to note that the control force is not active during escape; we assume that an individual prioritizes escape over attraction to its roost and minimization of its energetic costs *[9]*. Similarly, there are no aerodynamic forces in our model *[5]*. For our agents to qualitatively resemble the flying motion of birds, we visualize agents with delta-shaped bodies, the coloration of starlings, and enforce a `banking' motion, i.e., individuals roll according to their turning rate *[10]*.

The drag force $(\overrightarrow{\psi_i^f})$ keeps individuals around a predefined speed ('cruise speed', u), given that birds have a preferred speed from which they cannot deviate for long. Thus, 'drag' in our model is a spring force: the more an agent deviates from its cruise speed, the more it is pulled back to it (with weighting factor $w_u$). The drag force acts in both sturnoids and predoids.

*Motion integration – Steering vector*

The final vector that pushed agents into their next positions, referred to as 'steering vector' is calculated as the sum of the defined social forces and the locomotion forces. The drag force is incorporated in the integration time step, while all other forces are calculated every update step and remain constant during the integration timesteps until the next update. Note that the total information of all agents in the simulation is 'frozen' throughout a full update cycle and thus the sequence of individual updates doesn't alter the perceived environment.

At each update step of sturnoids the total update-force is calculated:

$$\overrightarrow{\psi_i^U} = \overrightarrow{\psi_i^{coord}} + \overrightarrow{\psi_i^w} + \overrightarrow{\psi_i^{control}} + \overrightarrow{\psi_i^{esc}} \tag{13}$$

If a sturnoid is in a flocking state, the escape force $\overrightarrow{\psi_i^{esc}}$ is 0. Similarly, if it is in an escape state, the control force $\overrightarrow{\psi_i^{control}}$ is 0. The update force of the predoid during pursuit is composed only by $\overrightarrow{\psi_p^h}$.

Finally, the total force applied at each integration step to all agents (sturnoids and predoid) is composed by:

$$\overrightarrow{\psi_i}(t) = \overrightarrow{\psi_i^U} + \overrightarrow{\psi_i^f}(t) \tag{14}$$

We then update the velocity and position of each agent following the midpoint method:

$$\vec{u_i}\left(t+\frac{dt}{2}\right) = \vec{u_i}(t) + \vec{a_i}(t)\frac{dt}{2} \tag{15}$$

$$\vec{r_i}(t+dt) = \vec{r_i}(t) + \vec{u_i}\left(t+\frac{dt}{2}\right)dt$$

$$\vec{a_i}(t+dt) = \frac{\overrightarrow{\psi_i}(t+dt)}{m_i}$$

$$\vec{u_i}(t+dt) = \vec{u_i}(t+\frac{dt}{2}) + \vec{a_i}(t+dt)\frac{dt}{2}$$

with *dt* being the integration step in our simulations.

*Initialization*

All sturnoids are initialized in a group formation; positioned within a sphere of a specific diameter with approximately the same direction and speed. This resembles the initial conditions of starlings when attacked by the RobotFalcon: the flock is cohesive and initiates flight towards the same direction, away from the robotic predator. The predoid is initialized in a random position and heading, far away from the flock. The first 5 seconds of the simulation are not included in the analysis to avoid any effect of the initialized density. The model does not use any input data.

*Parameterization and calibration*

The collective motion of sturnoids in the model is parameterized according to previous empirical findings *[4, 7, 11]*, insights from a previous model of collective motion of starlings *[5]*, and model calibration so the collective motion of starlings in the model resembles flocks of real starlings in certain properties. Unfortunately, individual trajectories of starlings from previous studies are not available and thus our model validation was based on distributions of nearest neighbor distances, speed, stability of neighbors, and velocity correlations *[4, 5, 11, 12]*. The configuration file that includes all parameters used for each simulation is copied to an exported data folder (by default named *'sim_data',* and in a subfolder defined in the parameter file as *data_folder*), along with all chosen output (see below) at the end of each simulation. All parameters involved in the calculation of the steering forces are constant across states. The parameter controlling the frequency of updating personal information about the nearest neighbors and the predator changes between normal and alarmed flocking, to resemble increased reaction frequency of prey under attack *[13]*.

The transition matrices controlling the switching states of agents during a simulation are parameterized based on 3 extremes for low, medium, and high alertness (or 'stress'). At each update step, an interpolator creates the matrix of each agent based on its level of stress. An example of the probably to transition from alarmed flocking to escape in relation to distance to the predator and a transition matrix across different stress level are given in supplementary Fig. S10 and S11. All parameters are given in supplementary Table S1.

*Output*

Observers structure is used to calculate a series of metrics of collective behavior at run time and export them as *.csv* files at the end of a simulation. Analyzing data of large flocks and calculating these metrics in other data-analysis focused software (e.g., *[14]*) is computationally challenging and expensive. Thus, this functionality enables the extensive future use of the model, also by experts without a computational background. Specifically, the model exports 6 types of datasets at a given sample frequency ($f_s$):

1. **Timeseries**: the 'raw' simulated data, timeseries of individual position (*posx, posy, posz*), heading (*dirx, diry, dirz*), speed, acceleration (*accelx, accely, accelz*), state, nearest neighbor distance squared (*nnd2*), relative position to the flock's center (*dist2fcent*, *dirX2fcent*, *dirY2fcent*, *dirZ2fcent*), and a metric indicating the distance of a sturnoid to the edge of a flock (*border*). Specifically, let $\Phi_i$ be the set of flock-members of focal individual $i$ within a 'probing distance' $d$ ($\Phi_i = \{ j, |r_j - r_i| < d \}$):
$$\varphi_i = \frac{1}{|\Phi_i|} \sum_{j \in \Phi_i} norm(r_j - r_i)$$
The closer $|\varphi_i|$ is to zero, the closer $r_i$ is to the centroid of the neighbors in $\Phi_i$, hence in the 'interior' of the probing volume. For our border metric, $d$ is infinite with $\Phi_i$ containing all flock mates.
2. **Flock**: timeseries of the flocks in the simulations, since splits may occur, with their respective sizes ($N$), ids, and volumes (calculated based on an object-oriented bounding box that includes all flock members, as in *[5]*).
3. **Diffusion**: how the internal structure of the flock changes over time (as in *[15]*). Specifically, this file contains the timeseries of global diffusion ($R$) and stability of neighbors ($Qm$) over a user-defined number of neighbors ($M$) and time-window *[11]*.
4. **Transition Matrices**: the timeseries of probabilities of switching to a different state given the current state of each sturnoid (*Pto0-Pto3*), along with its alertness (*stress*) and distance to the predoid (*dist2pred*).
5. **Information transfer**: the chain of copying a flockmate's escape maneuver, specifically the time of copying, the id of the individual copying a maneuver (*idx*) and of the one copying from (*src_idx*), and the copied state and substate (maneuver type, here 0 for turning and 1 for diving).
6. **Predator-prey**: information about the relative position of sturnoids and predoids.
7. **Correlation**: the timeseries of the correlation function of velocity fluctuations in relation to the linear size (L) of the flock, across different interaction ranges (as in *[12]*).

*Further functionalities*

A series of added functionalities are included in our model to support its future contribution to the study of complex patterns of starling flocks. Its interactive graphical user interface allows the monitoring of many individual variables such as the speed, nearest neighbor distance, state, and flock id in real time. In terms of escape reactions, in the simulations presented in the main text, a sturnoid selected which escape maneuver to perform stochastically, according to a parameterized probability. However, StarEscape also allows the probability of selecting a specific maneuver to depend on the local information of each sturnoid, such as its relative position to the predator or flock mates, or the predator's speed. For instance, when the predator is diving on the flock in high speed or the attacked individual is at the bottom of the flock, choosing a diving maneuver may be advantageous. Since empirical data to inform such individual reactions are not yet available, this functionality was not used in the present paper but could be further investigated over the model's parameter space. Other escape reaction, such as a 'zig-zag' maneuver (previously used to study agitation waves *[10]*) or extreme sudden turns (similar to individual behavior during flash expansions *[16]*) are also included in the model and can be added to a sturnoid's possible reactions or replace existing ones (level turn or dive).

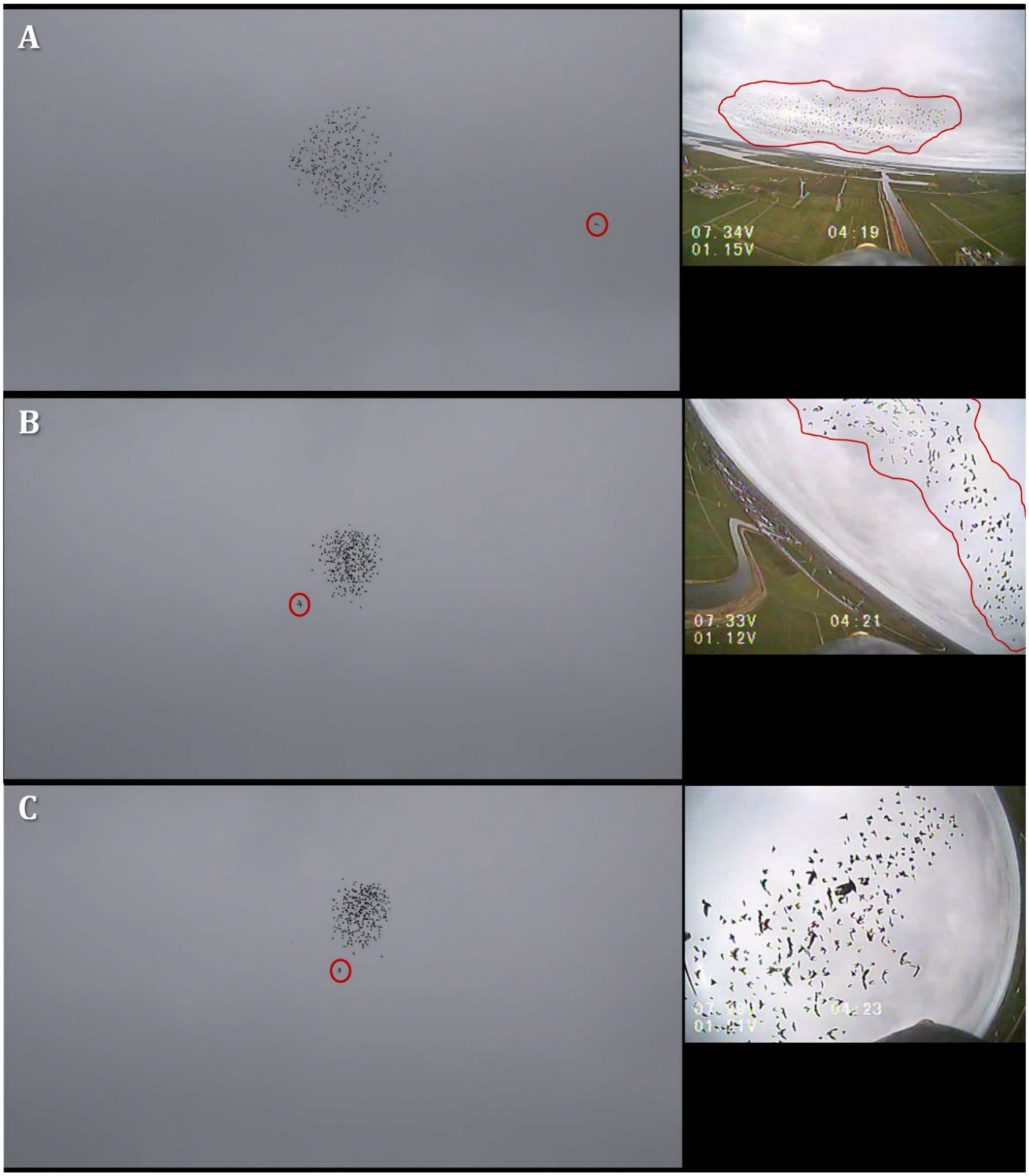


**Fig. S1. Screenshots from footage of the RobotFalcon pursuing a flock of starlings in the field**. The images on the left are from the ground camera and on the right from the camera on top of the robot. The red circles indicate the RobotFalcon's position in the ground camera frame, and the red freeform outlines the flock in the reference frame of the robot. A-C capture 3 consecutive instances of the pursuit (2 seconds apart).

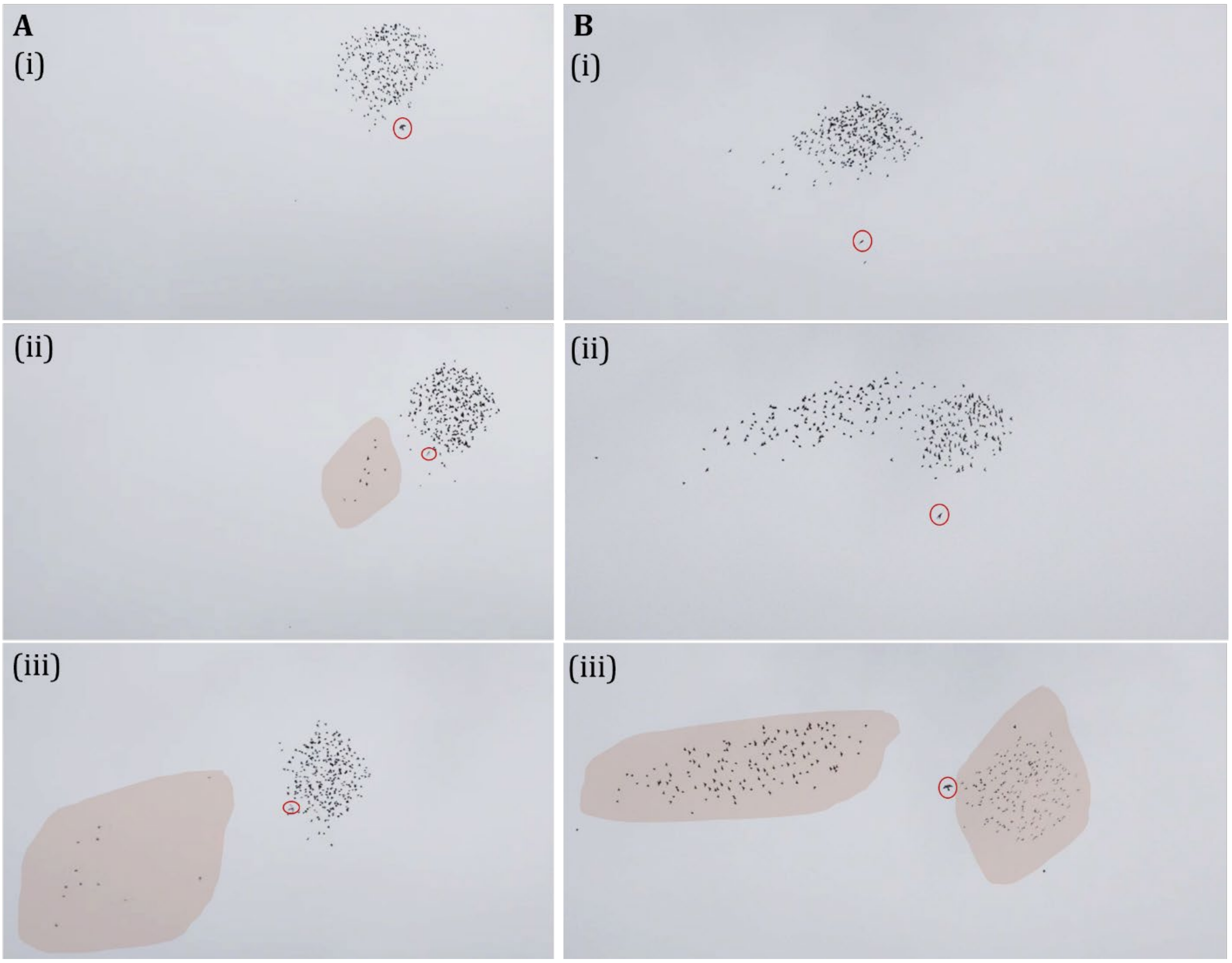


**Fig. S2. Splitting in footage of starlings pursued by the RobotFalcon (circled in red) in the field**. The shaded areas indicate the sub-flocks. **A.** A small loose sub-flock splits off from the main flock at the point of attack of the RobotFalcon. **B.** The flock splits into two sub-flocks by turning towards opposite direction in relation to the position of the RobotFalcon. The duration of each event (i-iii) was approximately 1 second.

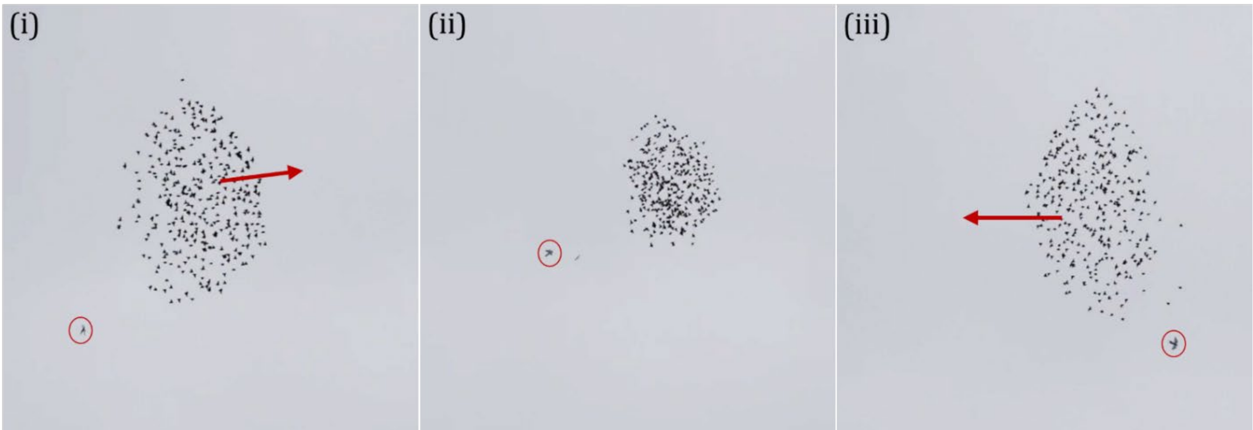


**Fig. S3. Compacting during collective turning** in footage of the RobotFalcon (circled in red) pursuing a flock of starlings in the field. The arrows indicate the direction of motion of the flock. The total duration of the screenshots (from i to iii) is approximately 3 seconds.

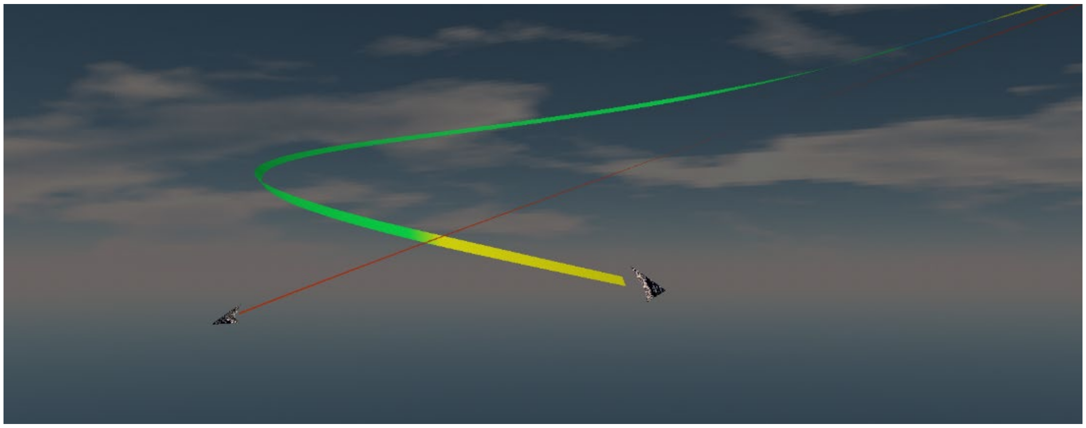

**Fig. S4. A sturnoid (green and yellow track) performing a diving-turn away from the predoid (red track).** This emerges by the individual performing a turn while recovering its pitch from a previous dive maneuver.

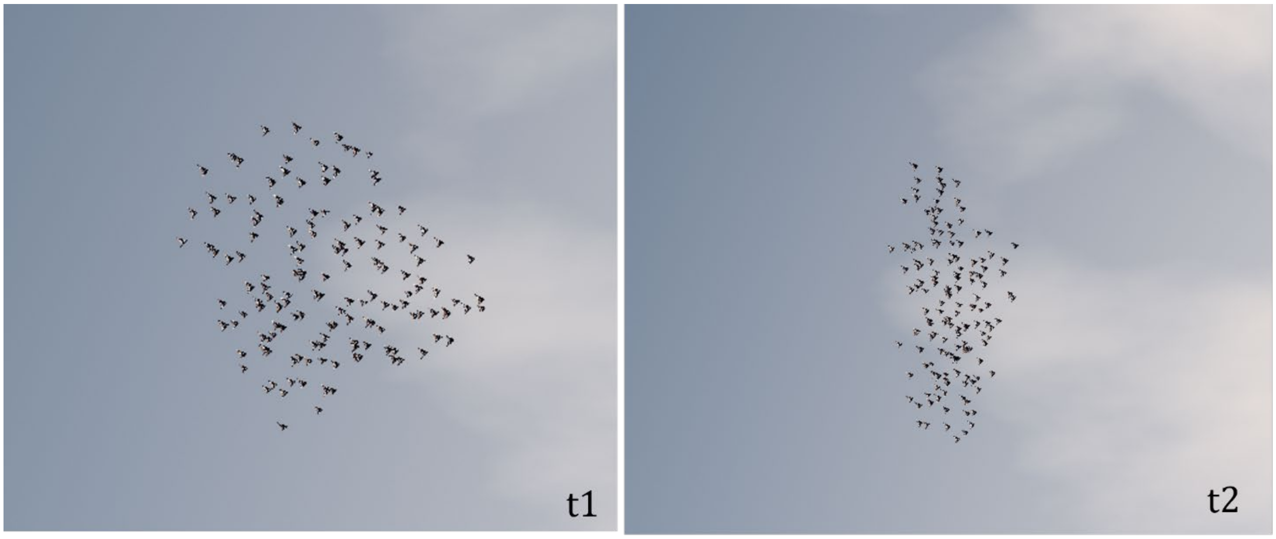


**Fig. S5.** The flock is compacting from t1 to t2as a result of an increased number of flock members being alarmed (higher reaction frequency) in the computational model StarEscape.

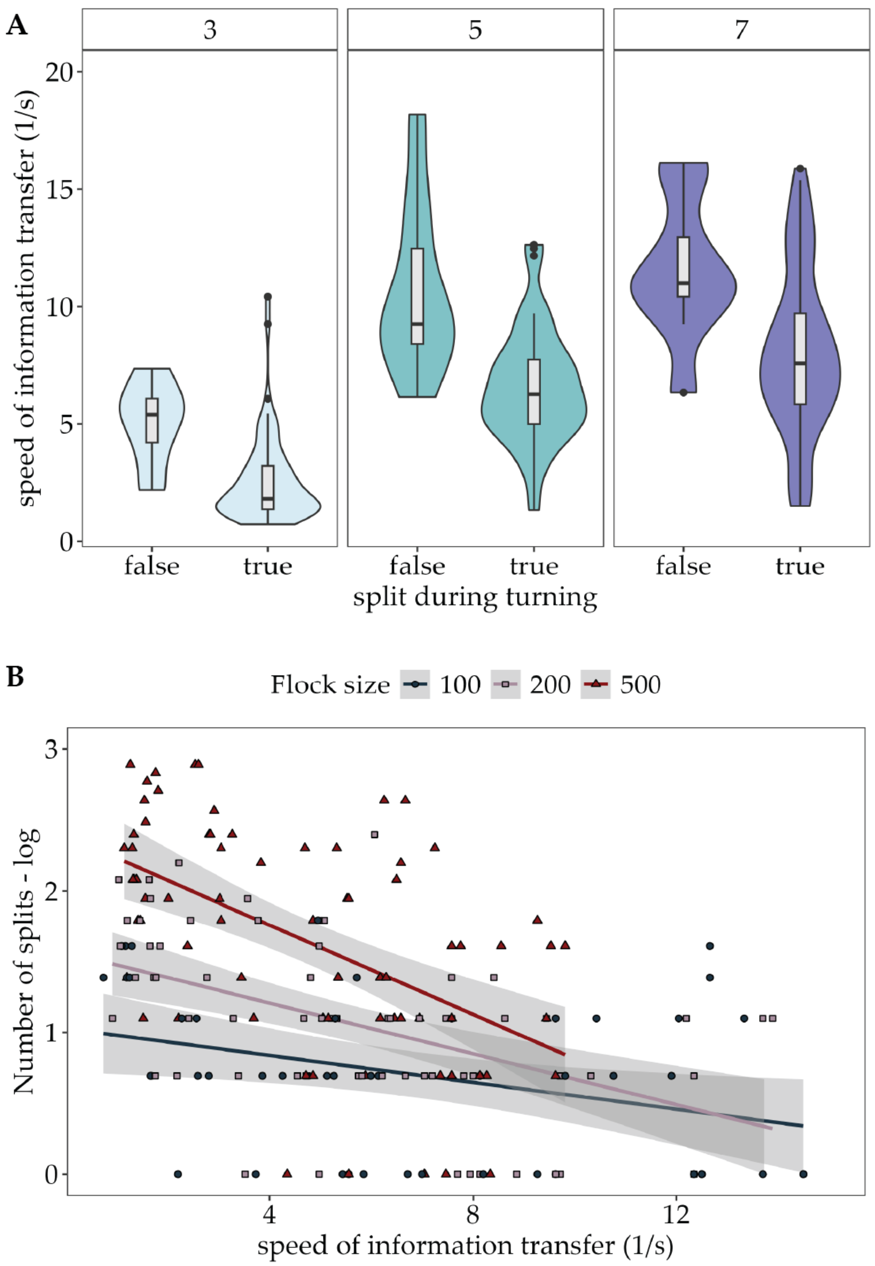


**Fig. S6. Information transfer and splitting during collective turns. A.** Slower information transfer (the inverse of the time for an escape maneuver to propagate through the flock) underlies the co-occurrence of splits and collective turns. Each panel corresponds to simulations with different topological range for copying an escape maneuver (parameter $n^{topo}{}_{\text{copy}}$; 3, 5 and 7 closest neighbors). **B.** The faster information propagates through the flock the less splits emerge. Each point represents a single flock during the first attack cycle of a simulation (first time the sturnoids react with a turning escape maneuver away from the predoid). A linear regression fitted to the logged number of splits (across all simulations with varying topological range for copying in which a split occurred; $R^2 = .42$, $p < .001$) shows the significant effect of speed of information transfer ($\beta = 0.96$, SE = 0.13, $p < .001$) and flock size ($\beta = 0.002$, SE = 0.0002, $p < .001$).

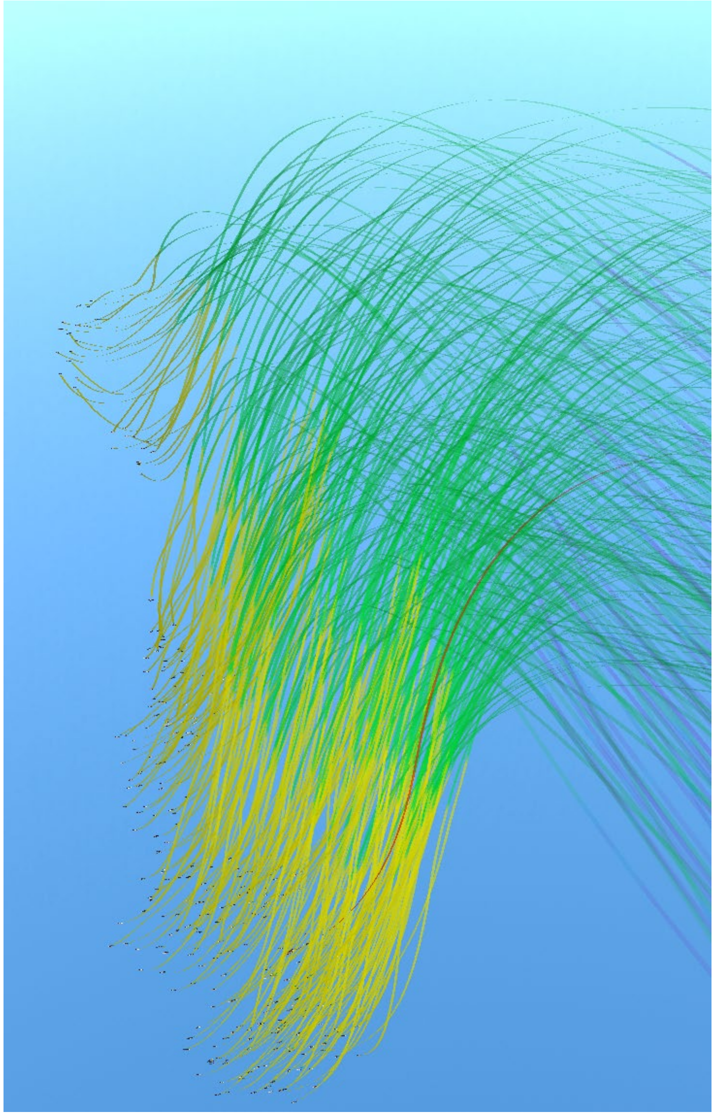

**Fig. S7. A flock of 300 individuals (green and yellow tracks) performing diving manoeuvres away from a predoid (red track).** The camera is positioned behind and on the side of the flock. The back part of the flock dives more (higher intensity) because they are closer to the predator. This results in the individuals that dive more ending up underneath the ones that dive less, which in turn leads to two main clusters. The green parts of the trajectories show the diving manoeuvre of each individual, while the yellow show each individual's recovery while returning its pitch to level.

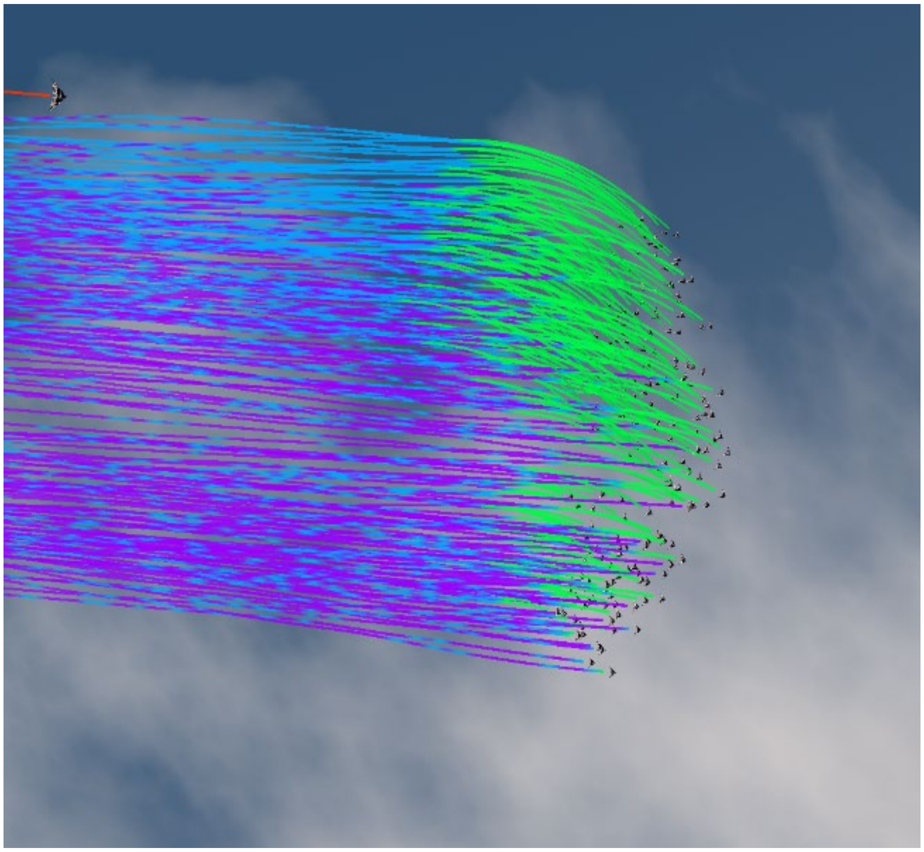

**Fig. S8. Propagation of an escape manoeuvre (level-turn) in a flock of 200 individuals, as a reaction to a sudden attack of the predoid (red track, top left).** The camera is positioned below the flock looking up. Sturnoids with purple tracks are in regular flocking, with blue in alarmed flocking, and with green are escaping. The individuals closer to the predator (top) have higher stress and thus transition to the alarm state and start turning away from the predoid first, to the left relative to their direction of motion and inwards in relation to the flock's centre.

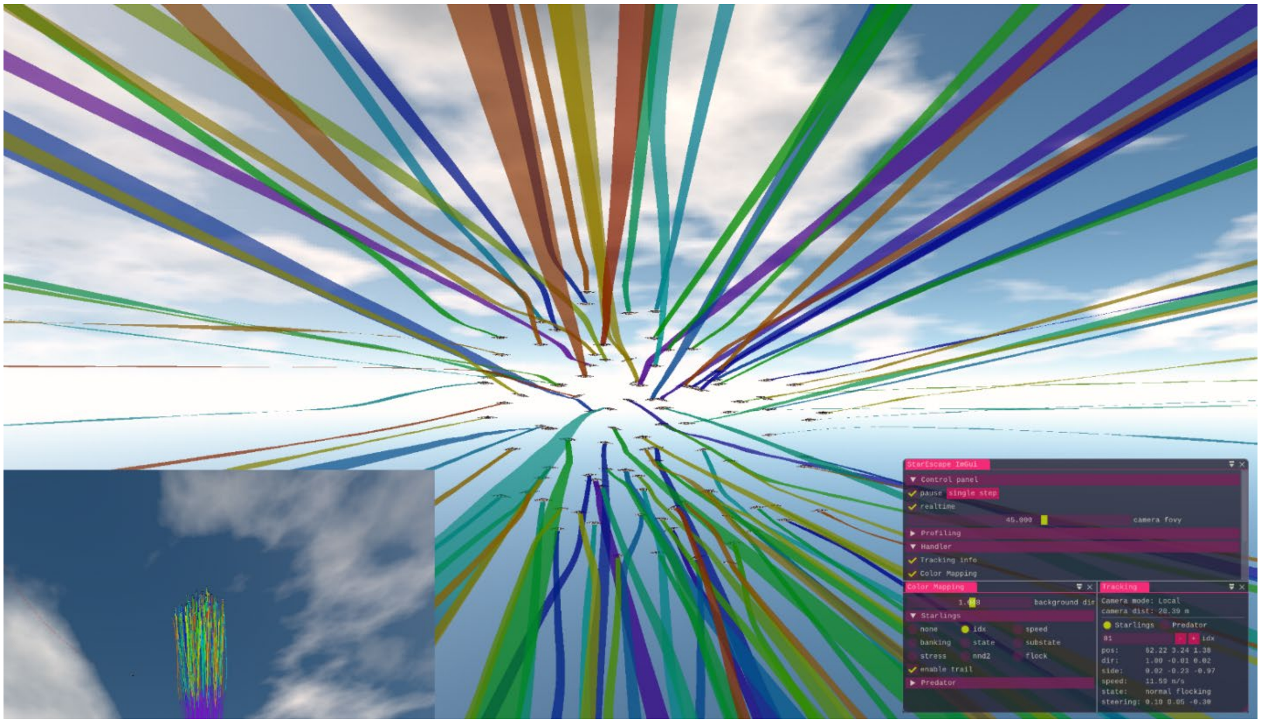


**Fig. S9. Simulation view** from the back of a flock (N=100), developed with OpenGL *[2]*. The left bottom panel shows a stable ground camera view following the flock from below and the right panel is the GUI that shows simulation specifics and controls the visualization (developed with DearImGui *[3]*). The tracks are coloured according to each individual's id (as seen also in the Color Mapping panel of the GUI).

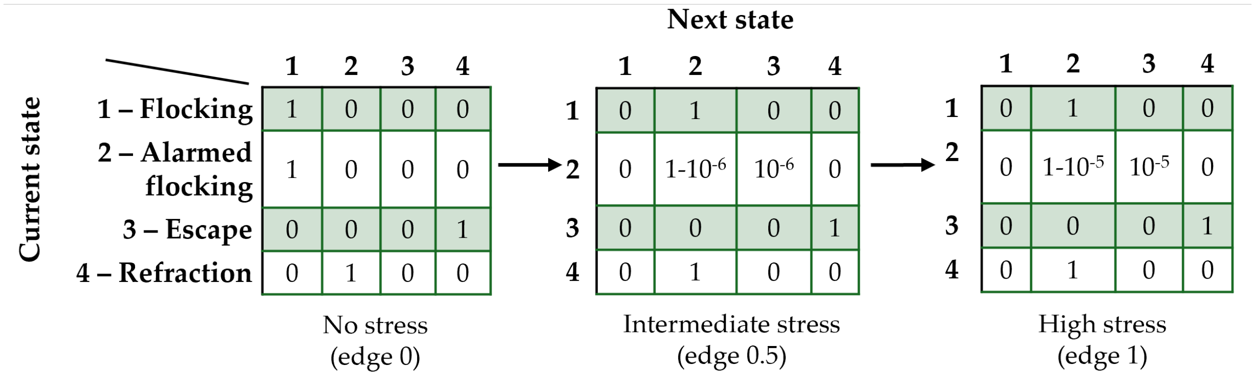


**Fig. S10. Parameterized transition matrices.** A linear interpolator creates a matrix for each agent based on the alertness level of the agent at each update step. The calculated probability (based on the current state of the agent) is then used to choose the next state of the agent. Note that states with a set duration (i.e., escape and refraction) use the matrix only after the duration of the state is finished ($t_{esc}$ and $t_{ref}$).

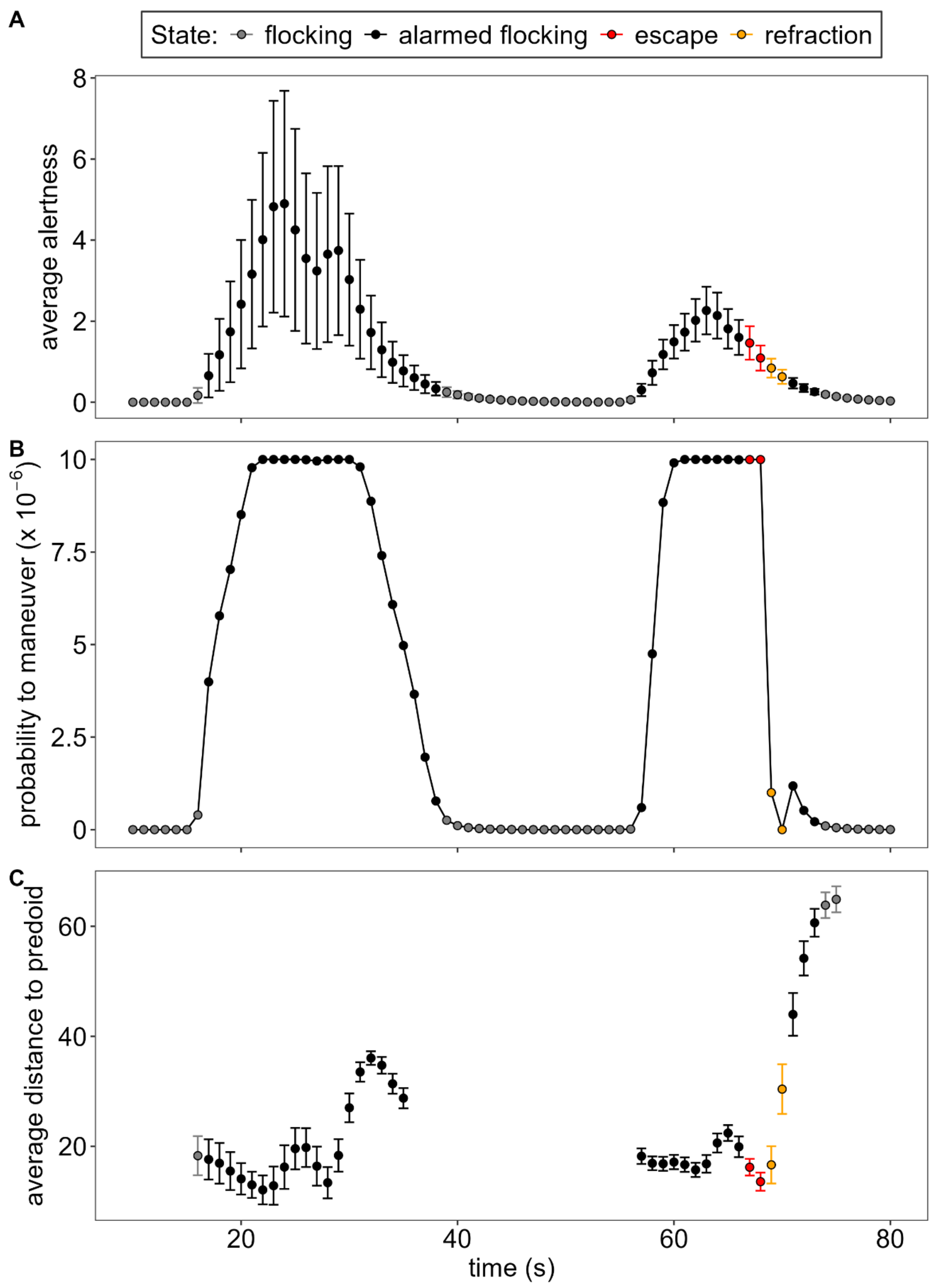


**Fig. S11.** Timeseries in a simulation with 100 individual of average **A.** alertness, **B.** probability to perform an escape maneuver (transition from alarmed state to escape) and **C.** distance to the predoid (with periods that the predoid is repositioned away from the flock omitted for visualization). The error bars indicate the standard deviation across all flock members.

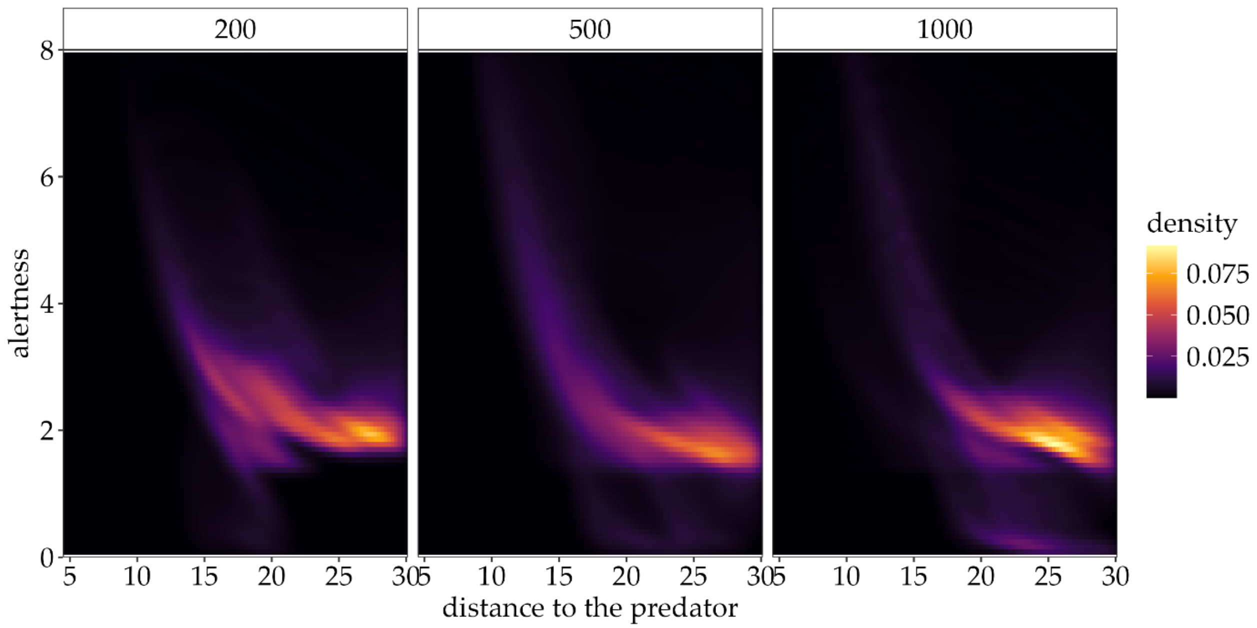


**Fig. S12.** Alertness (or stress) of sturnoids over distance to the predoid during alarmed flocking states in flocks of 200, 500, and 1000 individuals (within 30 meters from the predator).

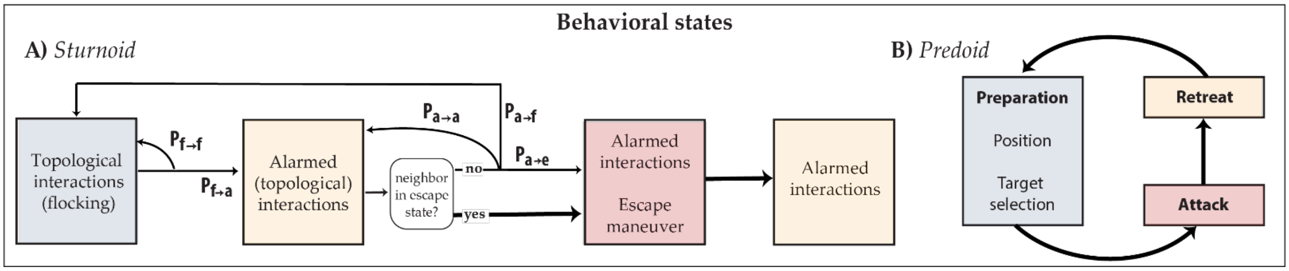


**Fig. S13. Behavioural states** and transitions of the starling-like (sturnoids) and predator-like (predoid) agents in StarEscape**.**

**Table S1. Parameters of the StarEscape model.** We calibrated our flocks to resemble cohesive flocks of starlings and performed a sensitivity analysis to ensure that small variation in these values does not result into changes in our model's performance. Parameters in the *Predation* section refer to the predoid, all others to sturnoids.

| Parameter | Name - Description | Value(s) |
|---|---|---|
| N | Flock size (sturnoids) | <=5000 |
| dt | Integration time step | 5 *ms* |
| $f_s$ | Sampling frequency | 0.1 *s* |
| ***Motion*** | | |
| u | Cruise speed | 9 *m/s* |
| $w_u$ | Weighting factor to return to cruise speed | 0.5 |
| $u_{min}$ | Minimum speed | *5 m/s* |
| $u_{max}$ | Maximum speed | 20 *m/s* |
| m | Body mass | 0.08 *kg* |
| $w_{alt}$ | Weighting factor of altitude attraction | 0.1 |
| $y_{alt}$ | Preferred altitude for attraction | 0 |
| $p_{alt}$ | Maximum pitch for altitude attraction | $45^{o}$ |
| $\sigma_{alt}$ | Smoothing parameter of altitude attraction | 200 *m* |
| $w_r$ | Weighting factor of roost attraction | 0.25 |
| | The x and z coordinates of the roost | [50, 100] |
| $\theta_r$ | Roost radius | 100 *m* |
| $w_n$ | Weighting factor of random noise (interval of uniform distribution) | 0.1 |
| ***Coordination*** | | |
| $\Delta t_r$ | Reaction time step - coordination | 50 *ms* |
| $\Delta t_{al}$ | Reaction time step - alarmed coordination | 25 *ms* |
| $\theta_{FoV}$ | Field of view | $270^{o}$ |
| $n^{topo}{}_{ali}$ | Topological range of alignment | 7 |
| $n^{topo}{}_{sep}$ | Topological range of avoidance | 1 |
| $n^{topo}{}_{coh}$ | Topological range of centroid attraction | 7 |
| $w_a$ | Weighting factor for alignment | 0.5 |
| $w_c$ | Weighting factor for centroid attraction | 1 |
| $\sigma^{l}{}_{coh}$ | Distance-based smoothing parameter for the weight of centroid attraction - min | 0 *m* |
| $\sigma^{h}{}_{coh}$ | Distance-based smoothing parameter for the weight of centroid attraction - max | 5 *m* |
| $w_s$ | Weighting factor of separation | 0.5 |
| $d_s$ | Minimum separation distance | 0.8 *m* |
| ***Escape*** | | |
| $n^{topo}{}_{copy}$ | Topological range of copying escape reaction | 7 |
| $t_{esc}$ | Duration of escape reaction | 3 *s* |

*Continued on next page*

*Table S1 – continued from previous page*

| | | |
|---|---|---|
| $t_{ref}$ | Duration of refractory period | 2 *s* |
| $\sigma_s$ | Parameter for the scaling of stress accumulation | 5 |
| $r_{decay}$ | Rate of decay of alertness | 1 $s^{-1}$ |
| $w_{stress}$ | Weighting factor of stress accumulation | 0.5 |
| $\theta_{esc}$ | Angle of level turn | $180^o \pm 5^o$ |
| $d_d$ | Maximum dive distance | 10 *m* |
| $\sigma_d$ | Smoothing parameter for dive maneuver | 10 *m* |
| $w_d$ | Weighting factor of diving | 0.5 |
| $P_{turn}$ | Probability to perform a turning manoeuvre | 0.5 |
| $P_{dive}$ | Probability to perform a diving manoeuvre | 0.5 |
| | High stress transition probability (max) from alarmed coordination to escape | 0.000001 |
| ***Predation*** | | |
| $d_{at}$ | Attack distance relative to the flock - level | 20 *m* |
| $\beta^l_{at}$ | Bearing angle to attack - level | $240^o$ |
| $\beta^a_{at}$ | Bearing angle to attack - altitude | $5^o$ |
| $t_{hunt}$ | Duration of attack | 60 *s* |
| $t_{retreat}$ | Duration of retreat | 30 *s* |
| $u_{scale}$ | Scaling of attack speed from the target's speed | 1.3 |
| $w_h$ | Weighting factor of attack | 10 |

#### Supplementary Movies

All movies are available on Figshare: https://doi.org/10.6084/m9.figshare.27606744.

**Movie S1.**
A flock of starlings pursued by the RobotFalcon. The following patterns of collective escape are observed in sequence or co-occurring (with the timestamp of occurrence in the Movie in seconds): blackening (upper part of flock, 00:00:10), columnar flocking (00:00:10), collective turn (00:00:11), collective turn (00:00:15), wave event (one pulse, downwards, 00:00:19), sub-flock split (bottom individuals, 00:00:21), collective turn (00:00:20), collective turn (00:00:24), cordon (00:00:25), blackening (sub-flock below the cordon, 00:00:26), collective turn (00:00:26), and split (in 2 flocks, 00:00:27). The simplified schematic representation of Fig. 1B is based on the patterns of collective escape observed here. The playback speed of the video is reduced to 40%.

**Movie S2.**
Individual escape maneuvers in starlings. **A.** A flock of starlings performs consecutive collective turns while being pursued by the RobotFalcon (large 'bird'). Individuals react to the artificial predator with level turns, some of their individual tracks are marked with colored lines (yellow, green, red, pink). **B.** A flock of starlings taking a columnar shape by individuals diving away from the RobotFalcon. The playback speed of the video is reduced to 40%. **C.** A flock splits, with the individuals closer to the predator and the point of splitting performing a different evasive maneuver: a level-turn (yellow and blue tracks) and a diving turn (red and pink tracks).

**Movie S3.**
A sequence of collective escape in a flock of 500 individuals in the model StarEscape. The color of each track in the ground view (inset, footage on the bottom left) shows the state of each individual, with purple being regular flocking, blue alarmed flocking, green escaping, and yellow alarmed flocking during refraction time. The flock performs two collective turns away from the predoid (red track), during which singletons and small sub-flocks split off. After the predoid is far away from the flock (00:00:15 s) and retreats (00:00:24 s), the flock dilutes as it returns to regular flocking.

**Movie S4.**
Recording of a simulation with StarEscape. A flock of 700 individuals first forms a columnar flock (as a result of diving manoeuvres) and subsequently a collective turn while in this formation, with some individuals diving further and a subgroup splitting off (see inset).

**Movie S5.**
Flock dilution in starlings a few seconds after the termination of the pursuit. At the beginning of the video, the RobotFalcon retreats.

**Movie S6.**
A close-up view of a small flock of 20 individuals performing consecutive collective turns away from the predoid in StarEscape, with the camera following the flock from behind. The body shape and coloration of the sturnoids is changed from the delta shape with starling coloration to another bird shape.

**Movie S7.**

A large simulated flock of 1000 individuals performing collective turns away from the predoid and splitting in StarEscape. After the pursuit, the camera is turned around by the user and the 3-dimensional shape of the flock is observed.
